# Metabolic and age-associated epigenetic barriers during direct reprogramming of mouse fibroblasts into induced cardiomyocytes

**DOI:** 10.1101/2024.02.28.582570

**Authors:** Francisco Santos, Magda Correia, Rafaela Dias, Bárbara Bola, Roberta Noberini, Rita S. Ferreira, Pedro Domingues, José Teixeira, Tiziana Bonaldi, Paulo J. Oliveira, Christian Bär, Bruno Bernardes de Jesus, Sandrina Nóbrega-Pereira

## Abstract

Heart disease is the leading cause of mortality in developed countries novel regenerative procedures are warranted for improving patients well fare. Direct cardiac conversion (DCC) can create induced cardiomyocytes (iCMs) and besides holding great promise, lacks clinical effectiveness for cardiac regeneration as metabolic and age-associated barriers remains elusive. Here, we identify by histone post-translational analysis that DCC triggers major age-dependent alterations in the epigenetic landscape. Metabolomics revealed decrease abundance of anabolic metabolites related to TCA cycle and glutaminolysis and profound mitochondrial network remodeling, with increased elongation and interconnectivity and reliance in mitochondrial respiration in iCMs.

Importantly, adult-derived iCMs present increase accumulation of oxidative stress in the mitochondria and pharmacological activation of mitophagy increase DCC of adult fibroblasts *in vitro*. Metabolic modulation *in vitro* and dietary manipulations *in vivo* improves DCC efficiency and are accompanied by significant alterations in the histone acetylation and methylation landscape and mitochondria homeostasis. Our study provides evidence that metaboloepigenetics as a direct role in cell fate transitions driving direct cardiac conversion into iCMs, highlighting the potential use of metabolic modulation in increasing cardiac regenerative strategies.

## INTRODUCTION

Cardiovascular diseases (CVDs) are the leading cause of mortality in developed countries. CVD-related pathologies are typically characterized by the irreversible loss of cardiomyocytes (CMs), which eventually leads to heart failure. Although conventional, mainly symptomatic treatments exist, novel regenerative procedures are warranted for improving cardiac regeneration and patients’ welfare.

The use of cellular reprogramming and transdifferentiation for regenerative biology and age-related pathologies is a promising area of research. Direct reprogramming of resident cardiac fibroblast (CFs) by forced expression of cardiogenic transcription factors (TFs) into induced cardiomyocytes (iCMs) has emerged as an attractive strategy (Ieda et al., 2010; Qian et al., 2012; Song et al., 2012; Tani et al., 2018). Despite holding great promise, transdifferentiation of resident adult CFs into iCMs still lack effectiveness (Hashimoto et al., 2018; Santos et al., 2020). One of the underlying barriers is aging, which imposes several epigenetic restrains to somatic cell reprogramming and cardiac regeneration (Bernardes de Jesus et al., 2018; Santos et al., 2020). In the past, we have identified a key molecular node that facilitates the reprogramming of aged fibroblasts safeguarding stem cell pluripotency during aging (Bernardes de Jesus et al., 2018; Nóbrega-Pereira & de Jesus, 2020). Understanding the impact of systemic barriers (including age-associated), responsible for limiting cell fate conversion, may offer new opportunities for cardiac regenerative strategies.

While following injury the capacity for regeneration of the adult mammalian heart is limited, the neonatal heart is capable of substantial regeneration; this capacity, however, is lost at postnatal stages (Costa et al., 2022; Soonpaa et al., 1996). Interestingly, this is accompanied by a shift in the metabolic pathways and energetic fuels preferentially used by CMs, switching from embryonic glucose-driven anaerobic glycolysis to adult oxidation of substrates, mainly fatty acids (FAs), in the mitochondria (Lehman & Kelly, 2002; Lopaschuk et al., 1992). Apart from bioenergetics, metabolites are key regulators of epigenetic events and gene expression programs driving stem and cancer cells fate decisions (Intlekofer & Finley, 2019; Lynch et al., 2020; Nóbrega-Pereira et al., 2018). In particular, systemic metabolic modulations, such as FA-depleted diets, inhibition of FA oxidation (FAO), or enhanced glucose metabolism, have been shown to boost postnatal CMs proliferation in mice and extend the regenerative window (Cardoso et al., 2020). The modulation of glucose and lipid metabolism can greatly impact the intracellular pools of acetyl-CoA and other metabolites, such as α-ketoglutarate (α-KG). Indeed, the availability of these substrates regulates the epigenetic landscape in pluripotent stem cells, and skeletal muscle stem cell specification and regeneration, through histone acetylation and methylation (van Gastel et al., 2020; Yucel et al., 2019).

Despite its importance, our understanding of the systemic metabolic and (aging-associated) epigenetic determinants in cell lineage commitment driving direct cardiac conversion (DCC) remains elusive. As dietary fats and hyperlipidemia are associated with CVDs, it is a priority to define the impact of systemic lipids in the transdifferentiation of fibroblasts into iCMs and cardiac regeneration. Here, we generated iCMs from mouse fibroblasts at different ages and subjected to differential lipid regimens to characterize the metabolic and histone acetylation signature of iCMs and the impact of systemic lipids and aging in DCC, allowing an unprecedent characterization of the molecular and metabolic circuitries limiting cellular conversion and providing ground for improved metabolic interventions that boost regenerative strategies in the heart.

## RESULTS

### Epigenetic transitions and direct reprogramming into iCMs differ with age

DCC is an attractive cardiac regenerative strategy, however, it is still an inefficient process as several epigenetic barriers (including age-associated) need to be overcome. We used mouse embryonic fibroblasts (MEFs), established from embryos of E13.5 gestating females, and adult skin ear-derived fibroblasts (AEFs) from adult (4-9 months) of C57 mice, and iCMs were generated by retroviral-induced MGT transduction, using pMx-puro-MGT or pBabe-puro (empty backbone, used as mock/control), followed by puromycin selection (Figure 1a). We observed that, at 11 days post-transduction (D11), retroviral-induced MGT transduction resulted in the increased expression of cardiac troponin T (cTnT) (Figure 1b-b’) and tropomyosin (Figure 1c) in iCMs, which was more evident in transdifferentiated MEFs compared to AEFs,highlighting the decreased efficiency of DCC with aging. During DCC, an early activation of cardiac genes (D3) takes place, which is followed by a deactivation of fibroblast genes at around D10 (Liu et al., 2016). qPCR analysis at D12 revealed increased expression of the reprograming factors in both MGT-transduced MEFs and AEFs (Figure 1d). However, induction of cardiac and repression of fibroblast markers (*Tnnt2* and *Col3a1*, respectively) was only detected in MGT-transduced MEFs (Figure 1d).

**FIGURE 1.**
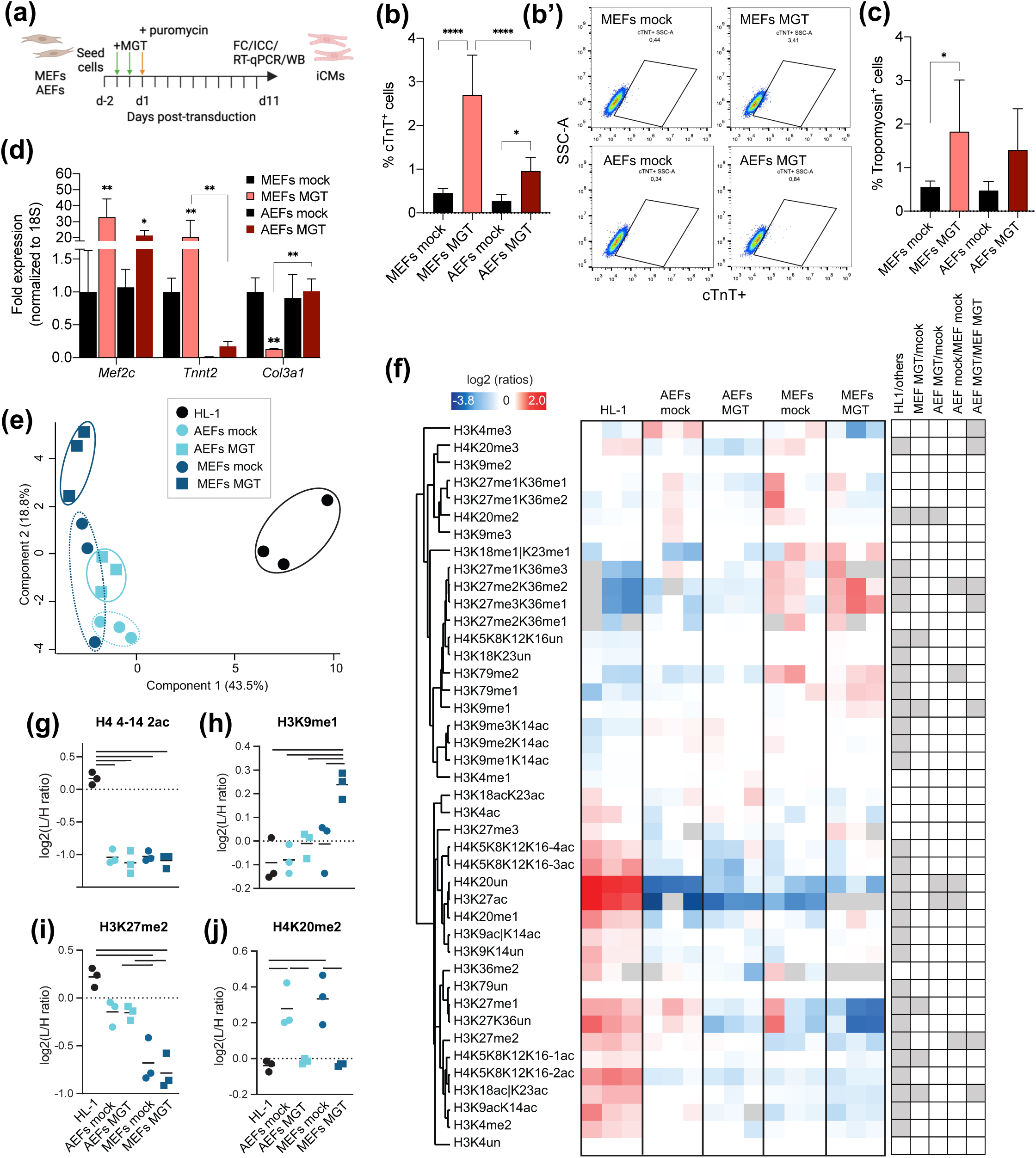
Epigenetic transitions and direct reprogramming into iCMs differ with age. (a) Schematic representation of the procedure for direct cardiac conversion of fibroblasts into iCMs. Fibroblasts (mouse embryonic, MEFs; adult skin ear fibroblasts, AEFs) are seeded followed by two rounds of MGT retroviral infections (green arrow, days −1 and 0) and puromycin selection at day 1 1 post-transduction (orange arrow). iCMs are analyzed 11 days post-transduction (D11) or the indicated days by several analytical techniques. (b-c) Flow cytometry quantification of cardiac troponin (cTnT) (b), representative histograms (b’) and expression of tropomyosin (c) in MEFs and AEFs mock or MGT retroviral-infected on D11/12. (d) qPCR analysis of the relative expression of the indicated genes in MEFs and AEFs mock or MGT retroviral-transduced on D11/12. (e-j) Mass spectrometry-based histone acetylation and methylation profiling in mock and MGT-transduced MEFs and AEFs on D11, and HL-1 mouse cardiomyocyte cell line. (e) Principal component analysis (PCA) based on histone PTM data obtained from the z scores of the samples shown in (f). Heatmap display of relative histone PTM levels. L/H (light/heavy) relative abundances ratios were obtained using a spike-in strategy (light channel: sample, heavy channel: spike-in standard), and were normalized over the average ratios across samples. Histone peptides were clustered based on Pearson’s correlation. The grey color indicates peptides that were not quantified. The panel on the right shows significant changes for the indicated comparisons (f). Display of selected histone modified peptides showing the levels of bi-acetylated form of H4 4–17 (g), mono-methylated form of H3 lysine 9 (h), di-methylated form of H3 lysine 27 (i) and mono-methylated form of H4 lysine 20 (j), the bar indicates a p-value<0.05. Graphical data are mean ± s.d. of n = 6-11 biological replicates from three independent experiments (b, c) or n = 3 biological replicates (d-j) from one representative experiment. Each point on the plot indicates individual measurements (e, g-j) and mean (g-j). p values were calculated by one-way ANOVA followed by post-hoc Tukey’s test. *p < 0.05; **p < 0.01; ***p < 0.001 with respect to mock-infected (d) or between the indicated groups. Graphs were created using GraphPad Prism (RRID:SCR_002798) and Perseus (RRID:SCR_015753), schema (a) with BioRender Premium and flow cytometry plots (b’) with FlowJo (RRID:SCR_008520).

Re-patterning of chromatin (such as H3K27me3 and H3K4me3) and DNA methylation have been shown to take place during conversion of neonatal fibroblasts into induced CMs (Liu et al., 2016). To obtain insights into whether the chromatin landscape transitions that take place during DCC differ with age, we performed mass spectrometry-based bulk analysis of histone post-translational modifications (PTMs, lysine acetylation and methylation marks) in transdifferentiated MEFs and AEFs, and corresponding mock-infected, at D11, along with the HL-1 mouse cardiomyocyte cell line (Figure 1e,g and Table S1). Principal component analysis (PCA, Figure 1e) based on histone PTMs data obtained from the z scores of the samples presented in the heatmap (Figure 1f) shows clustering of samples into three groups (HL-1, MEFs MGT and the remaining). A clear distinction can be observed between fibroblasts (regardless of age and transduction) and the HL-1 cardiomyocytes, with separation in the component 1. A less pronounced separation between transdifferentiated MEFs (MGT) and the remaining fibroblasts is observed in component 2 (Figure 1e), suggesting a remodeling of the chromatin in MEF-derived iCMs. Hierarchical clustering of differentially modified histone PTMs revealed clustering in two groups: a first cluster (21/42 total modifications) mainly composed of histone 3 (H3) methylated residues and H4K20me2, with increased levels in fibroblasts; and a second cluster (21/42 total modifications) with increased levels in HL-1 cells, which contains several histone H3 and H4 acetylated peptides (Figure 1f). These results suggest that, compared to fibroblasts, the chromatin in CMs is enriched in acetylated and deprived in methylated H3/H4 residues (Figure 1f, g). During reprogramming, an early rapid activation of the cardiac program and later progressive suppression of fibroblast fate at both epigenetic and transcriptional levels have been reported (Garry et al., 2021; Liu et al., 2016). Accordingly, we observe histone PTMs suggesting e a progression towards transcriptional silencing at D11, especially in MEFs, with DCC being accompanied by a progressive increase of marks typically associated with transcriptional repression, such as H3K27me3K36me1and, and decrease of H3K4me3, typically associate with active promoters (significant between AEFs MGT/MEFs MGT, Figure S1a, S1b). Other significant changes in histone PTMs were observed particularly in DCC of MEFs (MEFs mock/MGT) and between embryonic and adult iCMs (MEFs MGT/AEFs MGT). For instance, an increase in H3K9me1 (Figure 1h) and a decrease of H3K27me1 (Figure S1c) was observed in embryonic-derived iCMs only. Differences between adult and embryonic cells were also detected, with MEFs (mock and MGT) presenting lower levels of H3K27me2 (Figure 1i). A striking decrease in H4K20me2, was detected in both embryonic and adult iCMs, resembling HL-1 cardiomyocytes (Figure 1j). These results show significant epigenetic alterations with age in DCC, suggesting that substantial metabolic transitions take place during this process.

### Transdifferentiation of mouse fibroblasts into iCMs is accompanied by metabolic and bioenergetic transitions resembling cardiomyocytes

To understand in-depth the alteration in metabolic pathways taking place during transdifferentiation of fibroblasts into iCMs, we performed LC-MS-based metabolomics. PCA analysis (Figure 2a) shows clustering of samples into the three groups (HL-1, MEFs mock and MEFs MGT), with a clear separation of MEFs mock and MGT-transduced along Dim2, revealing that transdifferentiation of embryonic fibroblasts into iCMs is accompanied by pronounced alterations in the metabolome. A separation between the HL-1 cell line and MEFs along Dim1 is indicative of a clear distinction between mouse cardiomyocytes (HL-1) and MEFs, regardless of MGT transduction. Two-dimensional hierarchical clustering from univariate analysis of the 50 most significantly modulated metabolites (Figure 2b) revealed a primary split in the upper hierarchical dendrogram, with HL-1 cells clustering from MEFs, as denoted from the PCA separation. The samples cluster in two groups of relative abundance: a first smaller cluster of only 5 metabolites with higher relative abundance in HL-1 cells and a second group containing the remaining metabolites, with increased abundance in MEFs (Figure 2b). In both clusters, MEFs mock and MGT present significant differences in the abundance of metabolites, where, in some cases, MEFs MGT resemble HL-1 cells, as for increased phenylethylamine (Figure S1d), and decreased L-glutamine (Figure S1e) and adenine (Figure S1f). Differential analysis of cellular metabolites between mock and MGT-transduced MEFs was associated with 18 metabolic pathways constituted by the most differently expressed metabolites (*p*-value < *0.05*) (Figure 2c, S1g and Table S3) (Maurício et al., 2022). From this analysis, alanine, aspartate and glutamate metabolism (*p*-value < 0.0001); phenylalanine metabolism (*p*-value < 0.001); nicotinate and nicotinamide metabolism (*p*-value < 0.01); phenylalanine, tyrosine and tryptophan biosynthesis (*p*-value < 0.01) and taurine and hypotaurine metabolism (*p*-value < 0.05) represent the most significant altered pathways (Figure 2c and S1g). Top significantly altered pathways are related to the TCA cycle and anaplerotic reactions, including glutaminolysis, for the replenishment of precursors for amino acids and nucleic acids biosynthesis. Of note, MGT-transduced cells present a general decrease in relative abundance of key metabolites as compared to mock, including the alanine, aspartate and glutamate-related metabolites citrate, L-glutamate, succinate (Figure 2d-f) and L-glutamine (Figure S1e), whereas MGT-transduced cells present increased abundance of N-acetyl-L-aspartic acid (Figure 2g), a metabolite synthesized in the mitochondria from the amino acid aspartate and acetyl-CoA. As for phenylalanine metabolism, MGT-transduced MEFs present decreased abundance of L-phenylalanine and L-tyrosine (Figure 2h, i) and higher phenylethylamine as compared to mock-transduced MEFs (Figure S1d). The fetal metabolic shift is characterized by cardiomyocytés cell cycle-arrest and a shift from glycolysis to mitochondrial respiration and an associated cell cycle arrest (Lopaschuk et al., 1992). In agreement, cell cycle analysis by propidium iodide (PI) labelling shows that MEFs MGT exhibit an increase in G2/M arrest and a decreased proportion of cells in G0/G1 and S phase (Figure 2j). Interestingly, adult transdifferentiated fibroblasts show minor changes in the cell cycle (AEFs MGT versus mock) and, an expected, cell cycle arrest as compared to MEFs mock (decrease proportion in G0/G1, S and increase in G2/M, Figure 2j). The TCA cycle is a hub of metabolism, with central importance in both energy production and biosynthesis that can regulate cell fate transitions. Analysis of live TMRE staining, indicative of polarized mitochondria, reveals an overall increased intensity in embryonic compared to adult fibroblasts and a pronounced higher intensity in MEFs MGT at D11 (Figure 2k, k’). To get further insights into the bioenergetic switch during transdifferentiation of fibroblasts into iCMs, we measured the mitochondrial oxygen consumption rate (OCR) using Seahorse XF analyses (Figure 2l and Figure S1h). Both embryonic- and adult-derived iCMs present increased mitochondrial maximal respiration and spare respiratory capacity (Figure 2l), resembling the higher respiratory profile of HL-1 cells (Figure S1h) (McNally et al., 2021), which may reflect increased oxidative capacity, increased substrate provision or altered mitochondrial homeostasis during DCC. Transdifferentiation of embryonic fibroblasts into iCMs is accompanied by a unique increase in proton leak, and higher non-mitochondrial respiration is observed in adult-derived iCMs (Figure 2l). No major differences in the ECAR (Figure S1i, an indirect indication of glycolytic flux, or the overall energetic profile (Figure S1j), as the relative utilization of mitochondrial versus glycolytic pathways for energy production were observed. Overall, these data suggest that, in parallel to cell cycle arrest, a decreased engagement in anabolic pathways and increased reliance on the TCA cycle to fuel the mitochondrial respiratory chain, may be required during transdifferentiation of fibroblasts into iCMs.

**FIGURE 2.**
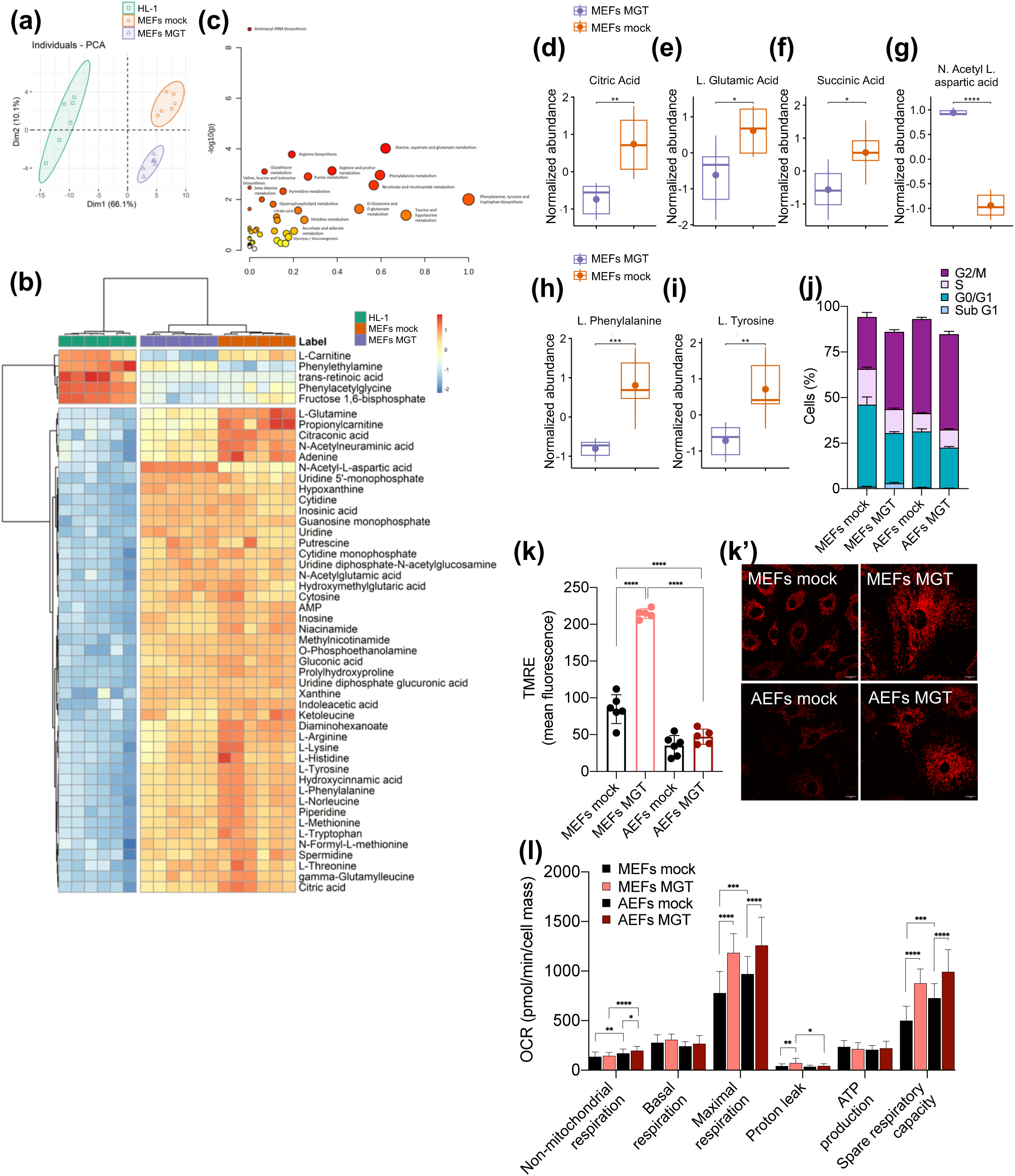
Transdifferentiation of mouse fibroblasts into iCMs is accompanied by metabolic and bioenergetic transitions resembling cardiomyocytes. (a-i) LC-MS-based untargeted metabolomics bulk analysis of MEFs mock or MGT-transduced on D11, and the HL-1 mouse cardiomyocyte cell line. PCA analysis score plot of the metabolite species dataset acquired for the indicated groups (a). Two-dimensional hierarchical clustering heatmap of the of the top 50 metabolites (p value < 0.05), comparing the 3 cell types based on relative abundance of each metabolite. At the top is the dendrogram of the samples, and on the left is the dendrogram of the identified species, statistical significance was determined by Kruskal–Wallis test (b). Dot plot of identified metabolomic pathways in MEFs mock and MGT on D11 associated with the indicated samples log(p) values from pathway enrichment analysis vs. values of the pathways impact (c). Pathway impact is based on enrichment analysis identifying the most relevant metabolic pathways and adjusted (log) p value, where higher impact values represent the relative importance of the pathway, the size of the circle indicates the impact of the pathway while the color represents the significance. Selected box-plot analysis relative abundances of identified metabolites citric acid (d), L-glutamic acid (e), succinic acid (f), N-acetyl-L-aspartic acid (g), L-phenylalanine (h), L-tyrosine (i) associated with alanine, aspartate and glutamate-related (d, e, f, g) and phenylalanine (h, i) comparison between MEFs mock (orange boxplots) and MEFs MGT (purple boxplots). (j) Cell cycle analysis by propidium iodide staining by flow cytometry displaying sum up bar graph of the percentage of cells in each phase of the cycle (sub-G1, G0/G1, S and G2/M) in MEFs and AEFs mock or MGT retroviral-transduced on D11. (k-k’) Confocal microscopy analysis of TMRE live cell imaging and quantification depicted as relative mean fluorescence intensity (k) in MEFs and AEFs mock or MGT retroviral-transduced at D13 and representative images (k’). (l) Cellular oxygen consumption rate (OCR) (non-mitochondrial-, basal-, maximal-, proton leak-and ATP-linked OCR, and spare respiratory capacity) in MEFs and AEFs mock or MGT retroviral-transduced at D11. Data were obtained using Seahorse XF96 Cell Mito Stress Test and normalized to cell mass using the sulforhodamine B (SRB) assay. Graphical data are mean ± s.d. of n=3-6 biological replicates from one representative experiment (a-i, j, k) or n=26-28 technical replicates from three independent experiments (l). Each point on the plot indicates individual measurements (a, k). p values were calculated by one-way ANOVA with False Discovery Rate multiple comparisons test by post-hoc Tukey’s test. *p < 0.05; **p < 0.01; ***p < 0.001; ****p < 0.0001 with respect to the indicated groups. Graphs were created using GraphPad Prism (RRID:SCR_002798), PCA graph (a), heat map (b) and box plots (d-i) were created using the R package pheatmap (RRID:SCR_016418) and R package ggplot2 (RRID:SCR_014601) and pathway analysis graph (c) with MetaboAnalyst 5.0 (RRID:SCR_015539). Scale bars, 20 µm in (k’).

### Extensive remodeling of mitochondrial network and mitophagy takes place during direct cardiac conversion into iCMs

Despite its importance, the extent of mitochondrial homeostasis and network remodeling taking place during transdifferentiation of fibroblasts from different ages into iCMs is largely uncharacterized. Immunofluorescence analysis of TOM20-labelled mitochondria, at D12, revealed increased total TOM20 fluorescence per cell in transdifferentiated MEFs, but not AEFs-derived iCMs (Figure 3a, b). No significant changes in the number of TOM20 particles were observed (Figure S2a), but an increase in the total area, indicative of size (Figure 3c) in both MEFs and AEFs-derived iCMs. DCC in MEFs is accompanied by significant remodeling of the mitochondria form factor, indicative of elongated shape (Figure 3d) and several parameters of mitochondrial network connectivity as branches per mitochondria, number and length in MGT-transduced MEFs only (Figure 3e and Figure S2b,d), resembling the HL-1 cells, with no significant alteration in AEFs-derived iCMs. During cardiomyocyte maturation, the increased reliance on mitochondrial respiration is accompanied by the production of reactive oxygen species (ROS) and adoption of FAs and pyruvate as main energy substrates (Lehman & Kelly, 2002). Indeed, adult-derived iCMs present higher lipid droplets, assessed by BODIPY 493/503 staining, compared to MGT-transduced MEFs, which may suggest a differential usage and mobilization of lipids (Figure S2e). Analysis of CellROX, indicative of cellular ROS (Figure 3f), revealed increased levels of oxidative stress during transdifferentiation of both embryonic and adult fibroblasts, with MEFs-derived iCMs presenting the highest levels, in agreement with proton leak measurements (Figure 2l), whereas increased MitoTracker red fluorescence, indicative of mitochondrial mass (Figure 3g), was only observed for MEFs-derived iCMs in line with previous observations (Figure 2k and Figure 3b-e). Strikingly, adult fibroblasts, both mock and MGT-transduced, present higher co-localization of ROS puncta in the mitochondria (Figure 3h, h’), suggestive of accumulated oxidative stress in the mitochondria, which may hamper transdifferentiation into iCMs. Mitochondrial network remodeling depends on processes such as mitochondrial biogenesis, mitophagy, and fusion and fission dynamics. No significant differences in the expression of Peroxisome proliferator-activated receptor gamma coactivator 1-alpha (PGC-1α), the master regulator of mitochondrial biogenesis, were observed upon transdifferentiation (Figure S2f, f’), Analysis of the autophagy marker LC3 revealed higher levels in adult compared to embryonic fibroblasts and increased in MEF-derived iCMs only (Figure 3i). Moreover, mitophagy was also defective during transdifferentiation of adult fibroblasts, as indicated by decreased LC3 puncta in the mitochondria compared to MEFs-derived iCMs (Figure 3j, j’). Consistent with this, MEF-derived iCMs present decreased p62 and increased PINK1, a ubiquitin ligase that initiates mitophagy (Lazarou et al., 2015), whereas less pronounced effects were detected for AEFs-derived iCMs (Figure 3k and Figure S2g). Next, we sought to determine if activation of mitophagy is required for iCM reprogramming. Previous reports suggest that autophagy is activated during early stage of iCM reprogramming in neonatal mouse cardiac fibroblasts (L. Wang et al., 2020). Consistent with that, treatment of MGT-transduced MEFs with rapamycin (10 nM) from D4 to D11, increased the percentage of cTnT+ cells in iCMs (Figure 3l and Figure S2h), with no major impact in AEFs-derived iCMs. Conversely, supplementation of MGT-infected AEFs with the mitophagy inducer urolithin A (UroA, 5 µM) (Ryu et al., 2016) increased the percentage of AEFs-MGT cTnT+ cells to the level of untreated MEFs-MGT (Figure 3l and Figure S2h, 2.26% vs 2.72%), whereas no major impact was produced in MGT-transduced MEFs. Of note, incubation with lysosomal inhibitor chloroquine (CQ) (25 µM) for 4 hours before cell harvest, blunted the effect of UroA in AEFs-MGT cells (Figure 3l). Overall, these results suggest that activation of mitophagy and subsequent clearance of damaged mitochondria, may be required for efficient transdifferentiation of fibroblasts into iCMs.

**FIGURE 3.**
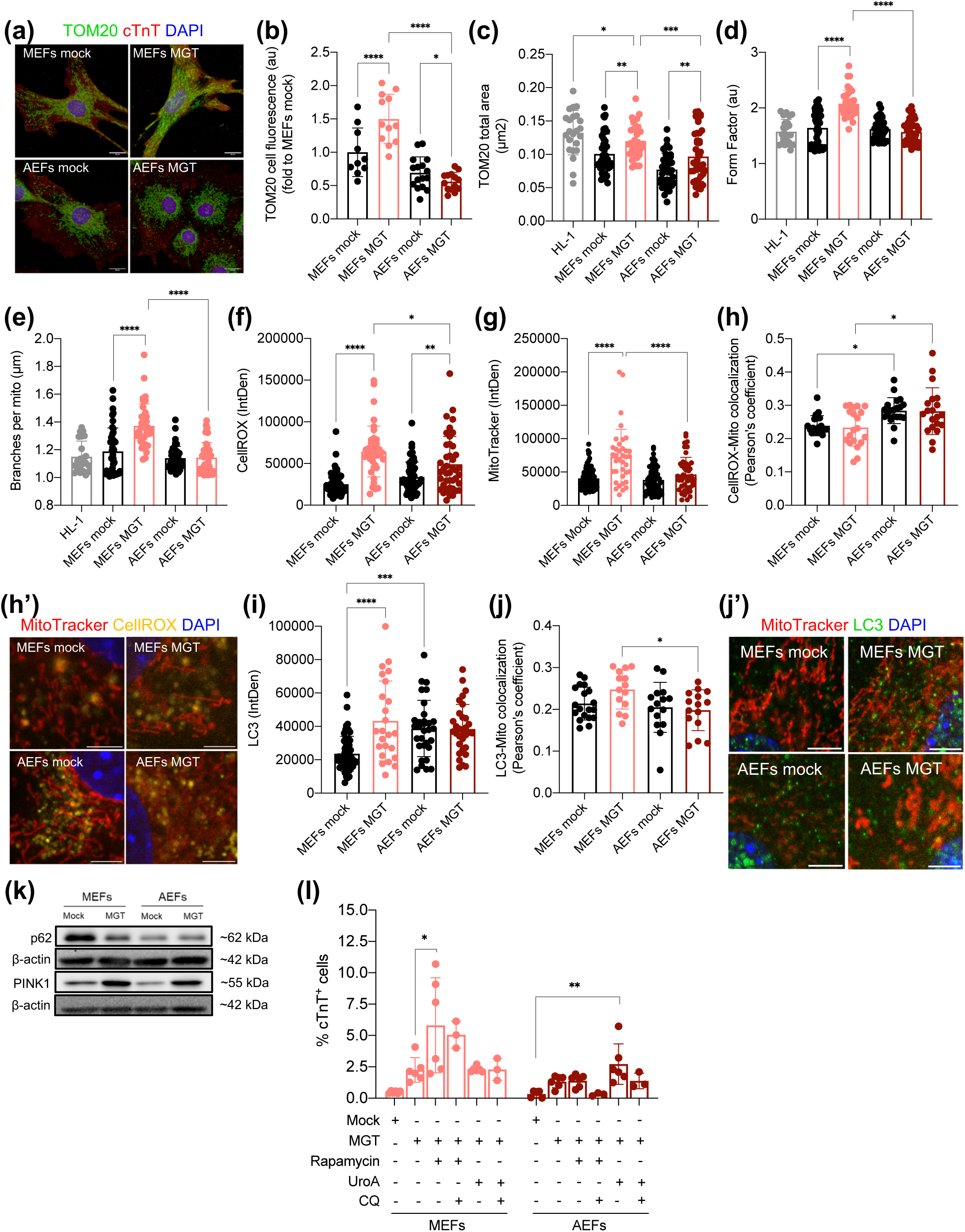
Extensive remodeling of mitochondrial network and mitophagy takes place during direct cardiac conversion into iCMs (a-e) Representative images of mitochondria TOM20 and cardiac troponin (cTNT) in confocal microscopy from MEFs and AEFs mock or MGT retroviral-transduced on D11/12, nuclei are labelled with DAPI (blue) (a). Analysis of total TOM20 fluorescence corrected by cell area (b), number of TOM20 particles (c), form factor (shape, d) and branches per mitochondria (network connectivity, e) in transdifferentiated MEFs and AEFs on D11/12 or HL-1 cells. (f-h’) Quantification of CellROX (ROS) (f) and MitoTracker red (g) per cell (integrated density, IntDen), and mitochondria-specific ROS puncta (Pearson’s coefficient) (h) and representative images of MitoTracker red and CellROX in confocal microscopy, nuclei are labelled with DAPI (blue) (h’) in MEFs and AEFs mock or MGT retroviral-transduced on D11. (i-j’) Analysis of LC3 signal per cell (IntDen) (i) and colocalization of LC3 to mitochondria (Pearson’s coefficient) (j) and representative images (j’) in MEFs and AEFs mock or MGT retroviral-transduced on D11 and representative images of MitoTracker red and LC3 protein (nuclei are labelled with DAPI, blue) in confocal microscopy, from MEFs and AEFs mock or MGT retroviral-transduced on D11. (k) Immunoblotting analysis for p62, PINK1 and β-actin in whole cell extracts from MEFs and AEFs mock or MGT retroviral-transduced on D11 (representative images, uncropped images of blots are shown in Figure S5). (l) Flow cytometry quantification of cardiac troponin (cTnT) in MEFs and AEFs mock or MGT retroviral-infected, at D11 untreated or supplemented with rapamycin (10 nM) or urolithin A (UroA, 5 µM) (treatment from D4 to D11). Chloroquine (CQ) (25 µM) was added for 4 hours before cell harvest in the indicated conditions. Graphical data are mean ± s.d. of n = 10/43 (a-e), n=20-83 (f-h), n=20-57 (i-j) cells analyzed or n = 6 biological replicates, from two independent experiments (l). Each point on the plot represents individual measurements (b-h, i, j, l). p values were calculated by Kruskal Wallis with false discovery rate correction and one-way ANOVA (b-e) and one-way ANOVA followed by Tukey’s post-test for multiple comparisons (f-j, l).*p < 0.05; **p < 0.01; ***p < 0.001; ****p < 0.0001 between the indicated groups. Graphs were created using GraphPad Prism (RRID:SCR_002798). Scale bars, 20 µm in (a), 5 µm in (h’ and j’).

### Metabolic modulation can bypass epigenetic and age-associated barriers to DCC

Mitochondria-derived acetyl-CoA and α-KG directly impact chromatin and promote an open poised chromatin state by contributing to histone acetylation and histone demethylation, respectively (Intlekofer & Finley, 2019). Importantly, pharmacological inhibition of histone-modifying enzymes that promote an open poised chromatin state have been shown to increase DCC efficiency (Fu et al., 2015; Hirai & Kikyo, 2014; Huang et al., 2018). To test the impact of mitochondria-derived metabolites in DCC efficiency, we took advantage of a transformed cell line derived from MEFs and harboring an α-myosin heavy chain (MHC)-eGFP reporter under the control of doxycycline (Dox) (Vaseghi et al., 2016), here referred as icMEFs. In the presence of Dox, expression of the MGT factors induce the cardiac fate easily observed by live GFP expression by flow cytometry and fluorescent microscopy (Figure 4a, a’ and Figure S3a). Importantly, upon Dox treatment for 3 days, we observed similar remodeling of mitochondria, as seen in transdifferentiated primary fibroblasts (Figure 3b, e) including increased mitochondrial total area, form factor and branches per mitochondrion (Figure S3b, S3d). Supplementation of Dox, together with glycolysis inhibitor 2-deoxy-D-glucose (2-DG, 1 mM) or in the presence of low glucose (LG: glucose 1mM, FBS 10%) or low lipids (LL: glucose 25 mM, FBS 1%) growth medium formulations increased the percentage of αMHC-eGFP-positive icMEFs and DCC efficiency compared to Dox only, as quantified by flow cytometry (Figure 4a, a’). Addition of Dox promoted an increase in mitochondrial mass in icMEFs as assessed by flow cytometry MitoTracker deep red staining (Figure 4b), similarly to MEF-derived iCMs (Figure 3g), and most of the manipulations were also accompanied by an increase in mitochondrial mass except for LL (Figure 4b). Supplementation of Dox produced an increase in ROS in icMEFs as detected by CellROX staining (Figure S3E), as previously observed in MEF-derived iCMs (Figure 3f). However, treatment with the cellular antioxidant N-acetylcysteine (NAC, 5 mM) had no impact in the percentage of αMHC-eGFP-positive icMEFs (Figure 4c). Also, despite the increase in mitochondrial mass produced by supplementation of resveratrol, a known activator of PGC-1α (RVT, 20 nM, Figure S3f) no significant changes in the percentage of αMHC-eGFP were observed (Figure 4c). These data suggest that forcing mitochondrial metabolism (including energy metabolism) can promote DCC by mechanisms that do not necessarily rely on increased mitochondrial mass. Moreover, manipulation of energetic fuels in the growth medium of mock or MGT-transduced primary embryonic and adult fibroblasts reveal that supplementation with sodium acetate (precursor of acetyl-CoA) and α-KG (*: sodium acetate 5 mM and α-KG 1.5 mM) under low lipids (LL*) or standard (S*: glucose 25 mM, FBS 10%) growth medium conditions increased the expression of cTnT in MEFs and AEFs, respectively (Figure 4d, e) whereas LG medium had no impact in the DCC of primary fibroblasts.

**FIGURE 4.**
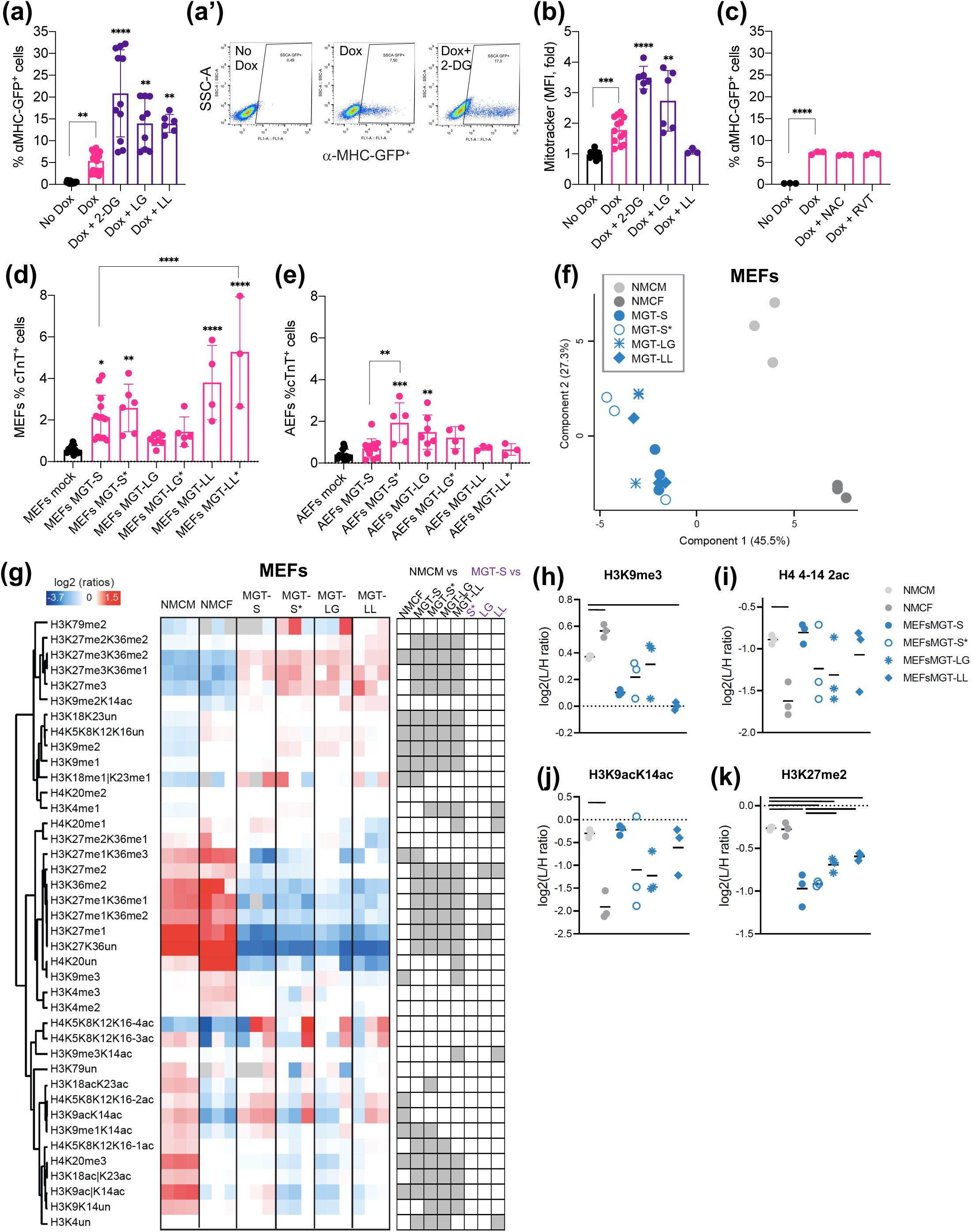
Metabolic modulation can bypass epigenetic and age-associated barriers to DCC. (a-a’) Flow cytometry quantification of the percentage of αMHC-eGFP (a) and representative scatter plots (a’) in icMEFs with no Dox or Dox for 3 days in the presence of 2-Deoxy-D-glucose (2-DG, 1 mM), low glucose (LG: glucose 1 mM, FBS 10%) or low lipids (LL: glucose 25 mM, FBS 1%) growth media formulation. (b) Flow cytometry quantification of mitochondrial mass by MitoTracker deep red staining depicted as relative median fluorescence intensity (MFI) in icMEFs with no Dox or Dox for 3 days in the presence of 2-DG (1mM), low glucose (LG: glucose 1 mM, FBS 10%) or low lipids (LL: glucose 25 mM, FBS 1%) growth media formulation. (c) Flow cytometry quantification of the percentage of αMHC-eGFP in icMEFs with no Dox or Dox for 3 days in the presence of N-acetylcysteine (NAC, 5 mM) or resveratrol (RVT, 20 nM). (d-e) Flow cytometry quantification of cardiac troponin (cTnT) in MEFs (d) and AEFs (e) mock or MGT retroviral-infected in the presence of standard (S: glucose 25 mM, FBS 10%), supplementation with sodium acetate and α-KG (S*: sodium acetate 5 mM and α-KG 1.5 mM), low glucose (LG: glucose 1 mM, FBS 10%) or low lipids (LL: glucose 25 mM, FBS 1%) alone or supplemented with sodium acetate and α-KG (LG*, LL*) growth media formulation, respectively, assayed on D12. (f-k) Mass spectrometry-based histone acetylation and methylation proteomic bulk analysis in neonatal mouse cardiac myocytes (NMCM) and fibroblasts (NMCF) and MEFs retroviral-infected on D11 in the presence of standard (S: glucose 25 mM, FBS 10%), supplementation with sodium acetate and α-KG (S*: sodium acetate 5 mM and α-KG 1 mM), low glucose (LG: glucose 1 mM, FBS 10%) or low lipids (LL: glucose 25 mM, FBS 1%) growth media formulation. PCA based on histone PTM data obtained from the z scores of the samples shown in heatmap display of histone PTM levels (f). L/H (light/heavy) relative abundances ratios were obtained using a spike-in strategy (light channel: sample, heavy channel: spike-in standard), and were normalized over the average ratios across samples (g). Histone peptides were clustered based on Pearson’s correlation. The grey color indicates peptides that were not quantified. The panel on the right shows significant changes between NMCM and MGT-S and the indicated comparisons. Display of selected histone modified peptides showing the levels of tri-methylated form of H3 lysine 9 (h), bi-acetylated form of H4 4–17 (i), mono-acetylated H3 lysine 9 and 14 (j) and di-methylated form of H3 lysine 27 (k), the bar indicates a p-value<0.05. Graphical data are mean ± s.d. of n = 3-16 biological replicates from two independent experiments (a, b, d, e) or n = 3 biological replicates from one representative experiment (c, f-k). Each point on the plot indicates individual measurements (a-f) and means (h-k). p values were calculated by one-way ANOVA followed by post-hoc Tukey’s test. *p < 0.05; **p < 0.01; ***p < 0.001; ****p < 0.0001 with respect to Dox (a, b), mock infected (d, e) or between the indicated groups. Graphs were created using GraphPad Prism (RRID:SCR_002798) and Perseus (RRID:SCR_015753) and flow cytometry plots (a’) with FlowJo (RRID:SCR_008520).

To get further insights as to whether the availability of mitochondria-derived metabolites (as acetyl-CoA and α-KG) can promote DCC by epigenetic changes, we performed histone PTMs in bulk analysis by mass spectrometry in transdifferentiated MEFs and AEFs under the different growth media at D11, and compared to the chromatin landscape of neonatal mouse cardiomyocytes (NMCM) and fibroblasts (NMCF) (Table S4). PCA shows clustering of samples into three groups: NMCF, NMCM and the remaining MEFs or AEFs conditions (Figure 4f and Figure S3g, respectively). A clear distinction can be observed between MEF-or AEF-derived samples and neonatal mouse cardiac cells (both cardiomyocytes and fibroblasts) in the component 1 (Figure 4f and Figure S3g, for MEFs and AEFs respectively) and a less pronounced separation between all mouse fibroblast cells (regardless of source, age and growth medium treatment) and cardiomyocytes (NMCM) is observed in component 2 (Figure 4f and Figure S3g, for MEFs and AEFs respectively). Despite the increased efficiency of DCC in MGT-transduced MEFs with supplementation of sodium acetate and α-KG (MGT-S*), low lipids (MGT-LL) or both (Figure 4d), no major differences in the chromatin landscape of iCMs generated under the different growth mediums was observed (Figure 4g). However, the level of certain histone residues in MEF-derived iCMs resembles NMCM and not NMCF, such as H3K9me3, H4K5K8K12K16-2ac, H3K9acK14ac (Figure 4h-j) and H3K9me1K14ac (Figure S3i) for iCMs generated under under standard conditions (MGT-S) and, to a minor extent, to MGT-S* and MGT-LL. Of note, iCMs generated under LL conditions (MGT-LL) present higher levels of H3K27me2 (Figure 4k) and H3K4un (unmodified) (Figure S3j) that differ from MGT-S and resembles NMCM. For AEF-derived iCMs, similar results were observed (Figure S3h). Similarly, the increased efficiency of DCC in MGT-transduced AEFs with supplementation of sodium acetate and α-KG (MGT-S*) (Figure 4e) is not accompanied by major differences in histone PTMs (Figure S3h). However, and as for MEF-derived iCMs, the increase in acetylated H4K5K8K12K16-2ac and H3K9acK14ac (Figure S3k and S3l), and the decrease in H3K9me1 (Figure S3m) for AEF-derived iCMs generated under standard (MGT-S) and sodium acetate and α-KG (MGT-S*) supplementation conditions, resembles NMCM and not NMCF. Overall, these results indicate that the acquisition of cardiac cell fate by embryonic and adult fibroblasts is accompanied by similar remodeling of the chromatin landscape, regardless of DCC efficiency instigated by growth medium manipulations.

### Dietary manipulation improves adult direct cardiac conversion into iCMs *ex vivo*

Dietary conditions have been shown to directly impact tissue histone acetylation levels and cell plasticity in cancer (Carrer et al., 2017; Pascual et al., 2021), however its impact in DCC remain elusive. We subjected young C57BL/6J adult male mice to several isocaloric dietary regimens including high fat (HFD, 60 kcal% fat), low fat/high fructose (LFD, 5 kcal% fat and 65% kcal fructose) and control purified ingredient (CD, 10 kcal% fat) diets for 10 weeks (Figure 5a). Animals subjected to HFD present a significant increase in body weight in respect to LFD and CD-fed animals (Figure S4a) and increase in glucose serum concentration (Figure S4b).

**FIGURE 5.**
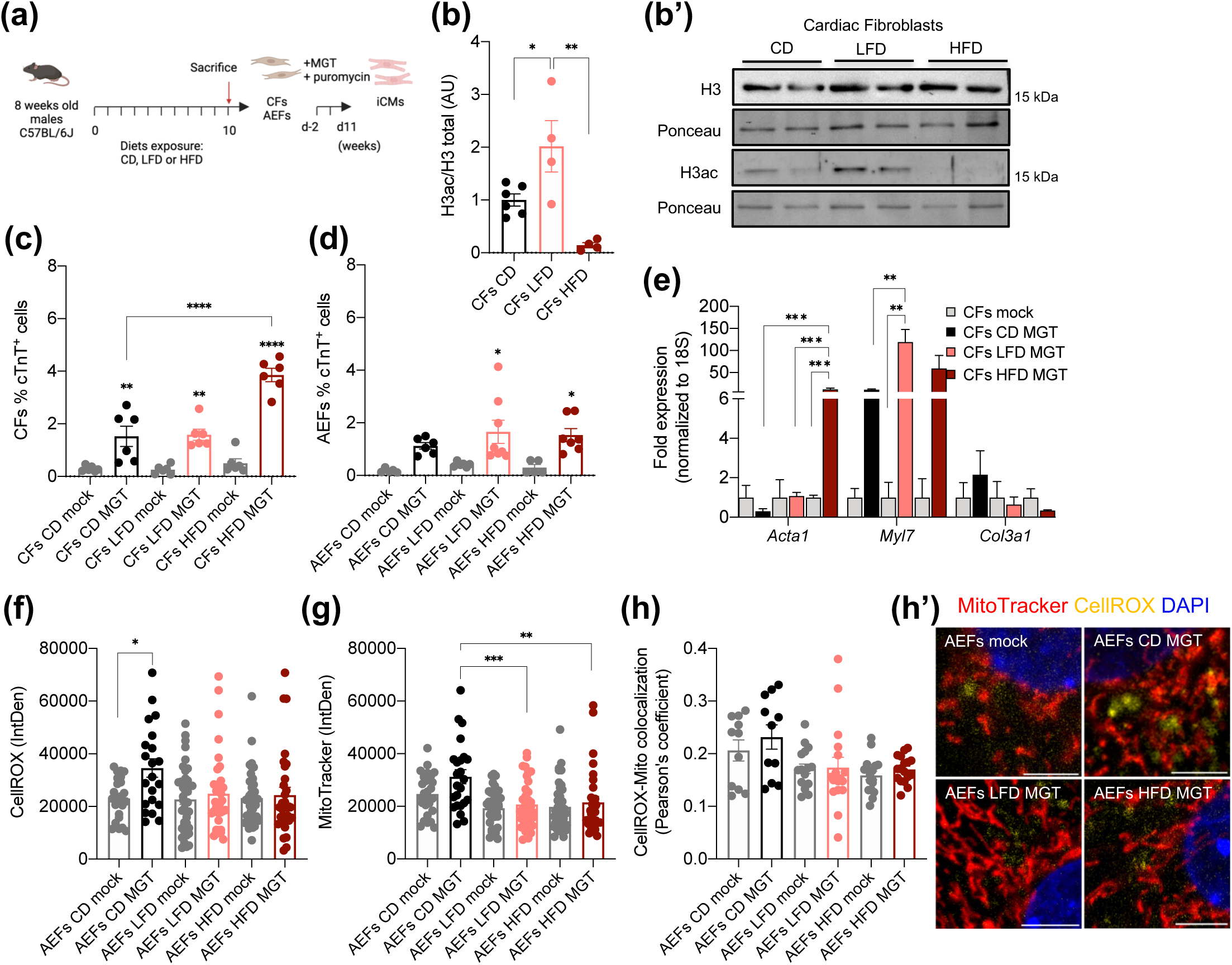
Dietary manipulation improves adult direct cardiac conversion into iCMs *ex vivo***. (**a) Schematic representation of the procedure. 8-week-old C57BL/6J adult male mice were subjected to high fat (HFD, 60 kcal% fat), low fat/high fructose (LFD, 5 kcal% fat and 65% kcal fructose) and control purified ingredient (CD, 10 kcal% fat) diets for 10 weeks. After sacrifice, cardiac and adult skin ear fibroblasts (CFs and AEFs, respectively) were isolated, seeded (D-2) and *ex vivo* MGT transduced followed by puromycin selection, and iCMs were analyzed on D11. (b-b’) Immunoblotting analysis of total histone H3 and H3 pan-acetylated displaying densitometric quantification of H3ac/H3 total (b) and representative images of histone H3, H3 pan-acetylated or Ponceau S staining (b’, uncropped images of blots are shown in Figure S5) from histone-enriched extracts of CFs isolated from animals under control diet (CD), high fat (HFD) and low fat/high fructose (LFD) at the end of the dietary regimen.(c-d) Flow cytometry quantification of cardiac troponin (cTnT) in CFs (c) and AEFs (d) mock or MGT retroviral-infected isolated from control diet (CD), high fat (HFD) and low fat/high fructose (LFD) at the end of the dietary regimen, assayed on D12. (e) qPCR analysis of the relative expression of the indicated genes in CFs isolated from animals under control diet (CD), high fat (HFD) and low fat/high fructose (LFD) at the end of the dietary regimen, and subjected to mock or MGT retroviral-transductions and analyzed on D11/12. (f-h’) Quantification of CellROX (ROS) (f) and MitoTracker red (g) per cell (integrated density, IntDen), and mitochondria-specific ROS puncta (Pearson’s coefficient) (h) and representative images, nuclei are labelled with DAPI (blue) (h’) in AEFs isolated from animals under control diet (CD), high fat (HFD) and low fat/high fructose (LFD) at the end of the dietary regimen, and subjected to mock or MGT retroviral-transductions and analyzed on D11/12. Graphical data are mean ± s.e.m of n = 4-8 biological replicates (b, c, d) or n = 11-24 cells analyzed, from two independent experiments (f-h), or n = 3 biological replicates from one representative experiment (e). Each point on the plot indicates individual measurements (b-d, f-h). p values were calculated by one-way ANOVA with False Discovery Rate followed by Tukey’s post-test for multiple comparisons. *p < 0.05; **p < 0.01; ***p < 0.001; ****p < 0.0001 with respect to the indicated groups. Graphs were created using GraphPad Prism (RRID:SCR_002798). Scale bars, 5 µm in (h’).

In order to determine whether exposure to different dietary regimens impacts histone acetylation and efficiency of DCC, we established *ex vivo* primary cultures of adult fibroblasts isolated from the heart (cardiac fibroblasts, CFs) and the skin (adult ear-derived fibroblasts, AEFs) from animals at the end of the dietary regimen (Figure 5a). Immunoblotting analysis revealed no differences in the levels of total histone H3 in CFs isolated from animals from the different diets (Figure 5b,b’ and Figure S4c), whereas, strikingly, an increase and pronounced decrease in H3 pan-acetylation (acetyl K9 + K14 + K18 + K23 + K27) was observed in CFs isolated from animals exposed to LFD and HFD, respectively (Figure 5b, b’ and Figure S4d), suggesting that nutritional stimuli can produce stable long-lasting histone PTMs (Carrer et al., 2017). Transdifferentiation of adult CFs into iCMs by MGT transduction revealed an increased expression of cTnT in CFs isolated from LFD and HFD mice, relative to mock-infected (Figure 5c) and higher DCC efficiency in MGT-transduced HFD CFs relative to CD (Figure 5c). Similarly, increased in cTnT expression by MGT transduction was observed in both LFD and HFD MGT-transduced AEFs compared to mock, contrasting the lower efficiency of CD-derived AEFs(Figure 5d), as previously observed. Accordingly, qPCR analysis revealed induction of the cardiac sarcomere structure genes *Acta1* and *Myl7*, and a tendency for repression of the fibroblast marker *Col3a1*, more pronounced in MGT-transduced CFs isolated from HFD and LFD than CD animals (Figure 5e). Similar expression levels of the reprogramming factor *Tbx5* were observed for all diet-isolated MGT-transduced AEFs (Figure S4e) and as for CFs, a tendency for more efficient repression of fibroblast markers (*Col1a1*, *Eln*) in LFD and HFD-isolated transdifferentiated AEFs (Figure S4e).

Diet-induced improvements in the transdifferentiation efficiency of (cardiac and skin) adult fibroblasts could be related to alterations in mitochondrial metabolism, as previously observed (Figure 2 and 3). No major difference in TMRE live staining for polarized mitochondria was observed in MGT-transduced AEFs between the dietary regimens (Figure S4f). Immunofluorescence analysis revealed higher ROS levels in transdifferentiated AEFs from CD animals (Figure 5f) and a tendency of lower cellular ROS levels in MGT-transduced AEFs isolated from LFD and HFD animals (Figure 5f). MitoTracker red staining (Figure 5g) revealed reduced mitochondrial mass in AEF-derived iCMs isolated from LFD and HFD as compared to CD animals, and a tendency to lower oxidative stress in the mitochondria (Figure 5h, h’), as previously observed for more efficiently generated embryonic derived-iCMs (Figure 3h, h’). This data suggests that decreased oxidative stress in the mitochondria of AEF-derived iCMs obtained from animals exposed to LFD and HFD may facilitate DCC.

## DISCUSSION

Aging has been reported to be a barrier for cell reprogramming (Nóbrega-Pereira & de Jesus, 2020), possibly due to abnormal chromatin states, or even the abundance and function of histone-modifying enzymes (Pal & Tyler, 2016). DCC is accompanied by significant changes in the epigenetic landscape, including re-patterning of the deposition of H3K4me3 and H3K27me3 (Liu et al., 2016). In particular, H3K4me3 has been shown to be promptly placed at the promoters of cardiac genes, while those on fibroblast-specific genes are removed at a slower rate (Liu et al., 2016) and the decreased levels of H3K4me3 observed here possibly reflect its removal from fibroblast-specific *loci*. Our results highlight substantial histone modifications during DCC, especially in more efficiently generated embryonic-derived iCMs, characterized by increased levels of H3K27me3K36me1, H3K9me1 and H3K27me1 (Figure 1f and S1). Contradictory roles have been attributed to the manipulation of H3K9me1 in DCC, either impairing or enhancing reprogramming, at early and later time points, respectively (Hirai & Kikyo, 2014; Ifkovits et al., 2014). H3K27me1 is associated with active transcription and correlates with the deposition of H3K36me3 in a Setd2-dependent manner (Ferrari et al., 2014). Remarkably, a decrease in H4K20me2 was observed in both MEF-and AEF-derived iCMs, resembling HL-1 cardiomyocytes (Figure 1j). This modification is enriched at sites of DNA damage and lower levels of H4K20me2 have been linked to decreased cell cycle progression (Paquin & Howlett, 2018). Indeed, during the fetal metabolic switch, mitochondrial fatty acid utilization increase chromatin oxidative stress and contribute for cell cycle arrest and CMs maturation (Menendez-Montes et al., 2021). iCMs are enriched in cell-cycle inactive genes, and blocking or synchronizing the cell cycle have been shown to improve DCC efficiency (Liu et al., 2017). MS-based metabolomics revealed significant alterations in the TCA cycle and glutaminolysis, including the anaplerotic replenishment of precursors for the biosynthesis of amino acids and nucleic acids (Figure 2b), suggestive of increased reliance on TCA cycle fueling mitochondrial respiration, further confirmed by higher maximal respiration and spare respiratory capacity in iCMs (Figure 2l). We further show that embryonic-derived iCMs display branched and complex mitochondrial network, which also takes place during CMs maturation (Lopaschuk et al., 1992). Remodeling of mitochondrial morphology and dynamics regulates several mitochondrial functions, as bioenergetics, apoptosis and ROS production (Hong et al., 2022). We show that MEF-derived iCMs present increased levels of ROS-labeling dyes and proton leak (Figure 2l and 3f). ROS (such as superoxide and hydrogen peroxide) act as secondary messengers playing important roles in cell fate regulation including cell reprogramming (Maryanovich & Gross, 2013; H. Wang et al., 2021). However, exacerbated oxidative stress can be detrimental for cellular homeostasis. We observe increased oxidative stress in the mitochondria of adult-derived iCMs (Figure 3h), suggestive of accumulated damaged mitochondria (Hong et al., 2022), which may hinder the maturation of the mitochondrial network and DCC efficiency. Moreover, compared to embryonic, adult-derived iCMs present decreased markers for activated mitophagy(Figure 3j, k’) and supplementation of the mitophagy-inducer UroA increased DCC efficiency in adult-cells (Figure 3l). Autophagy has previously been shown to be activated during early stage of DCC (H. Wang et al., 2021) and PINK1-Parkin-mediated mitophagy is essential during fetal maturation of CMs (Gong et al., 2015), preceding a boost in mitochondrial biogenesis (Persad & Lopaschuk, 2022). We found no evidence for alterations in mitochondrial biogenesis during DCC of embryonic or adult fibroblasts at the time-points studied (Figure S2a, f).

Several mitochondria-derived metabolites (including from the TCA cycle) serve as co-factors or substrates for epigenetic-modifying enzymes, linking metabolism to epigenetics and transcriptional regulation (Intlekofer & Finley, 2019). The histone PTM and metabolic reprogramming taking place during DCC led us to hypothesize that nutrient modulation may influence the availability of key metabolites that impact cell fate transitions. Indeed, inhibition of glycolysis or deprivation of lipids alone or supplemented with sodium acetate (which is converted into acetyl-CoA by ACSS2) and α-KG, consistently increased DCC *in vitro* in MGT-transduced primary cells and the icMEFs cell line and, moreover, adult skin fibroblasts derived from LFD-exposed animals present increased DCC (Figure 5d). MS-based histone proteomics revealed similarities between iCMs generated under low lipid conditions (MGT-LL) and NMCM, as higher levels of H3K27me2 (Figure 4k) and H3K4un (Figure S3j), suggesting that lipid deprivation increases DCC efficiency by direct remodeling of the chromatin status. This data is consistent with previous studies in pluripotent cells, where low exogenous lipids promote *de novo* lipogenesis and increase the contribution of acetyl-CoA to histone acetylation (Cornacchia et al., 2019), whereas high levels of α-KG promote histone and DNA demethylation to maintain pluripotency (Carey et al., 2015). Pharmacological manipulations that promote histone acetylation and demethylation (the so-called open-poised chromatin state) increase DCC efficiency *in vitro* (Singh et al., 2020; H. Wang et al., 2021). We observe that histone H3 acetylation levels are decreased and increased, respectively, in cardiac fibroblasts isolated from HFD-and LFD-fed animals (Figure 5b), as observed by others (Carrer et al., 2017). Both nutritional manipulations promoted increased DCC efficiency *ex vivo* (Figure 5c,d), reinforcing that complex chromatin transitions, in addition to histone pan-acetylation, regulate cell fate transitions in DCC. Remarkably, both LFD-and HFD-derived adult iCMs present decreased levels of oxidative stress in the mitochondria and reduction of mitochondrial mass (Figure 5), highlighting that improved mitochondrial clearance is a determinant factor for DCC efficiency in adult fibroblasts.

Overall, our study reveals that the availability of mitochondria-derived metabolites favor chromatin landscape transitions driving DCC, highlighting a dynamic cross-talk between mitochondrial homeostasis and epigenetics during cell state conversions. Moreover, we show that enhanced clearance of damaged mitochondria contribute for mitochondrial network maturation and function, as engagement in respiration, leading to improved direct cardiac conversion and the future design of more efficient heart regenerative strategies.

## EXPERIMENTAL PROCEDURES

### Mice studies

Male mice C57BL/6J (wild type, WT) were purchased from the Charles River Laboratories. All animal studies were performed at the iBiMED animal facility at University of Aveiro, reviewed and approved by the Ethical Commission of Department of Medical Sciences’ Animal Welfare Body (CEDCM-BEA) and licensed by the national regulatory agency Direcção Geral de Alimentação e Veterinária (DGAV). Mice were housed and handled in accordance with standard protocols of the European Animal Welfare Legislation, Directive 2010/63/EU (European Commission, 2016). 7-week-old male C57BL/6J mice were allowed to acclimatize for 1 week prior to initiation of the study. The mice were subjected to three dietary regimens including high fat (HFD, 60 kcal% fat D12492i), low fat/high fructose (LFD, 5 kcal% fat and 65% kcal fructose, D22060205i) and control purified ingredient (CD, 10 kcal% fat, D12450Ki, all from Research Diets) diets for 10 weeks (n=8-9 animals per diet, n=26 animals in total), using weight-matched littermate mice. The body weight was recorded on a weekly basis. Mice were sacrificed by cervical dislocation. At necropsy, blood was collected by cardiac puncture into centrifuge tubes, centrifuged for 15 min at 1600g and serum was stored at −80°C until analysis. The heart was dissected and skin tissue samples from the ear were collected and kept in PBS 1x at 4°C before fibroblast isolation.

### Determination of glucose level in serum

Glucose levels from serum samples of mice subjected to the different diets was measured using the Liquick Cor-GLUCOSE 60 Kit (Cormay Diagnostics 2-201) and following the manufacturer’s instructions. Absorbances were read at 500 nm, using a Tecan infinite M200 (Tecan) plate reader. The glucose concentration is defined by the ratio between the test absorbance and the standard absorbance, multiplied by the concentration of the standard.

### Isolation of mouse primary fibroblasts and cardiomyocytes

Primary fibroblasts were isolated as previously reported (Nóbrega-Pereira & de Jesus, 2020). Briefly, mouse embryonic fibroblasts (MEFs) were prepared from E13.5 embryos, whereas adult ear-derived fibroblasts (AEFs) were isolated either from adult male (4-9 months) or 18-week-old animals at the end of the CD, HFD, LFD dietary regimens. Isolation of neonatal (day 1-3) mouse cardiomyocytes (NMCMs) and cardiac fibroblasts (NMCFs) were performed according to (Lu et al., 2022). Briefly, homogenized hearts were transferred to a gentleMACS™ C Tube (Miltenyi Biotec), and washed twice with PBS 1x. The enzyme mix from Neonatal Heart Dissociation Kit mouse and rat (Miltenyi Biotec) was added to the tube and placed into the gentleMACS™ Dissociator (Miltenyi Biotec), for 1 hour. Growth medium consisting of Minimum essential medium (MEM, Bioconcept), supplemented with 5 % FBS (Gibco), 1% penicillin-streptomycin (Life Technologies) was added to the tube and the content was transferred to a falcon through a 70 µm cell strainer, and centrifuged at 300 g for 5 min. The pellets were resuspended in 10 mL of growth medium, pre-plated onto a 10 cm dish and incubated for 2 hours at 37°C with 1% CO_2_. For NMCMs plating, plates were previously coated with 0.02% of porcine gelatin, and cardiomyocytes present in the supernatant of the pre-plating mix were counted and plated with growth medium at 37°C with 1% CO_2_. For NMCFs, a pre-plating dish was washed 3 times and DMEM supplemented with 10% FBS (Gibco) was added to the adhered cells and incubated at 37°C with 5% CO_2_.

For isolation of adult mouse cardiac fibroblasts from 18-week-old male mice at the end of the dietary regimens (CD, HFD, LFD), mouse hearts were isolated and washed with ice-cold HBSS 1x with 1 % penicillin-streptomycin (Gibco). Hearts were cut into halves, and 3 hearts were pooled into a 10 cm dish. Hearts were washed with ice-cold HBSS containing 1% penicillin-streptomycin (Gibco) for 5 min to remove blood. The tissue was minced with a scalpel, mixed with of digestion buffer [Collagenase II (Worthington)/ Dnase I (SIGMA)/0.05% Trypsin-EDTA in 12 mL of HBSS] and incubated for 15 min at 37°C with 5% CO_2_. A 5 mL-pipette was used to dissociate the tissue by pipetting up and down, the dishes were again incubated for another 15 min at 37°C with 5% CO_2_. Up to 4 mL of supernatant with cloudy solution was removed, transferred to a 15 mL falcon and placed on ice. 4 mL of digestion buffer was added, and the remaining tissue clumps were dissociated with 1 mL pipette tip and incubated for 15 min at 37°C CO_2_. The digested cell solution was passed through a 100 µm cell strainer and centrifuged at 100 g for 2 min. The supernatant was collected and centrifuged at 300 g for 10 min. The supernatant was discarded slowly, and the pellet was resuspended in Fibroblast Growth Medium [FGM-3 (Promocell), supplemented with 10% FBS (Gibco), 1% penicillin-streptomycin (Gibco), insulin and FGF (Promocell)]. The isolated fibroblasts were seeded onto 10 cm dishes previously coated with 0.1% of porcine gelatin.

### Culture of mammalian cell lines and mouse primary fibroblasts

Human Embryonic Kidney 293T (HEK 293T) were purchased from ATCC between 2012 and 2016, and mouse HL-1 (Claycomb et al., 1998) and icMEFs (Vaseghi et al., 2016) were kindly provided by Dr. Christian Bär and Dr. Li Qian, respectively. All lines were tested as being mycoplasma-free and authenticated by examination of morphology and consistent *in vitro* performance. Cell lines and primary fibroblasts were cultured in DMEM (Gibco) high glucose (4.5 g/L glucose) or low glucose (1 g/L glucose), supplemented with 10% (v/v) heat-inactivated FBS (Gibco), 1 mM sodium pyruvate and 1% antibiotic-antimycotic (Gibco) at 37°C and 5% CO_2_ atmosphere. HL-1 cells were cultured in Claycomb medium (SIGMA) with 10% FBS, 1% penicillin-streptomycin (Gibco), 0.1 mM norepinephrine (SIGMA) and 2 mM L-glutamine (Gibco). icMEFs and iCMs (from day 1 post-transduction onwards) were cultured in DMEM/M199 (Zenbio) (4:1) with 10% FBS, 1 mM sodium pyruvate and 1% antibiotic-antimycotic.

### Direct cardiac conversion of fibroblasts into induced cardiomyocytes (iCMs) and cell treatments

For direct cardiac conversion, fibroblasts were retroviral-infected according to (L. Wang et al., 2015). Briefly, on day 1, 5×10^6^ HEK 293T cells were seeded onto a 10 cm dish. 4 μg plasmid DNA, either pMX-puro-MGT (Addgene, #111809), a polycistronic vector that expresses the reprogramming factors Mef2c, Gata4 and Tbx5 (MGT) or pBabe-puro (an emptyback bone vector, used as mock/control; Addgene, #1784) were mixed with 4 μg pCL-Ampho (retroviral packaging plasmid), together with 576 μL Opti-MEM (Gibco) media containing 24 μL of X-tremeGENE 9 DNA (Roche) and added to HEK cells. Medium was changed the next day (day 2) and fibroblasts were seeded at a density of 1.6×10^5^ cells per well in 6-well plates or 1.2×10^6^ cells in p100 plates coated with 0.1% gelatin (SIGMA). Virus containing-medium was collected 48 hours after transfection (day 3), followed by filtration through a 0.45 μm polyethersulfone filter and supplementation with polybrene (Millipore, 10 μg/mL), and added to target cells. This step was performed twice a day for 2 days. On day 5 (day 1 post-transduction; D1), the virus-containing medium was replaced with growth medium and changed every 3-4 days until analysis (D11/12). For positive selection, puromycin (SIGMA, 2 μg/mL) was added to target cells on D1 and maintained at a concentration of 1 μg/mL from D4 to D11.

For growth medium manipulation, mock or MGT retroviral-infected fibroblasts growth medium was replaced on D4 (and maintained until analysis on D11) by standard (S: glucose 25 mM, FBS 10%), supplementation with sodium acetate and α −KG (S*: sodium acetate 5 mM and α-KG 1.5 mM), low glucose (LG: glucose 1 mM, FBS 10%) or low lipids (LL: glucose 25 mM, FBS 1%) alone or supplemented with sodium acetate and α-KG (LG*, LL*). Supplementation with rapamycin (Tocris Bioscience, 10 nM) or urolithin A (UroA, SIGMA, 5 µM) was added on D4 until analysis on D11. Chloroquine (CQ, SIGMA 25 µM) was added 4 hours before cell harvest, on D11.

### Direct cardiac conversion using the icMEF cell line and cell treatments

icMEFs were seeded at a density of 1.5×10^4^ cells per well in 24-well plates coated with 0.01% gelatin. The following day (day 0), growth medium was either replaced with medium supplemented with doxycycline (SIGMA, 2 μg/mL) and drugs (2-deoxy-d-glucose (2-DG, SIGMA, 1 mM), N-acetylcysteine (NAC, 5 mM), resveratrol (RVT, 20 nM) or substituted by low glucose (LG: glucose 1 mM, FBS 10%) or low lipids (LL: glucose 25 mM, FBS 1%), both supplemented with doxycycline 2 μg/mL. On day 1 in the afternoon, supplemented growth medium was replenished. icMEFs were treated with dox, drugs and growth medium formulations for a total of 3 days.

### Flow cytometry assays

For live staining, cells were incubated with BODIPY 493/503 (ThermoFisher, 0,2 μg/mL) for 10 min at RT, MitoTracker Deep Red (Invitrogen M22426, 2 nM) for 15 min at RT, or CellROX Deep Red (ThermoFisher C10422, 2 μM) for 20 min at 37°C, Cells were dissociated with 0.025% Trypsin-EDTA, washed, and resuspended in PBS. For determining reprogramming efficiency in icMEFs, αMHC-eGFP was detected. For intracellular staining, iCMs on D11-12 were harvested by trypsin digestion for 5 min at 37°C. Cells were washed with ice-cold FACS buffer (DPBS supplemented with 2% FBS and 2 mM EDTA) and subjected to fixation and staining with Cell Fixation and Permeabilization Kits (BD Biosciences). Incubation with antibodies was performed for 1h at RT. The following antibodies were used: primary anti-mouse cardiac Troponin T (Invitrogen, MA5-12960, 1:200), anti-mouse Tropomyosin Alexa 488 conjugated (Santa Cruz Biotechnology, 1:200), goat anti-mouse Alexa 488 (Invitrogen A11001, 1:600). Cells were analyzed on a BD Accuri™ C6 Flow Cytometer (BD Biosciences) and FlowJo software (RRID:SCR_008520).

### Cell Cycle Analysis by PI Staining

Cells were collected and washed with PBS 1x complemented with FBS (2%) and centrifuged at 300 g for 5 min at RT. Cold ethanol (70%) was added dropwise followed by vortexing. Cells were fixed for 1h at 4°C, washed and resuspended in the Propidium Iodide Staining solution (SIGMA, 2 μg/mL) with RNAse A (1000 μg/mL, Thermo Scientific) and incubated overnight at 4°C. Cells were analyzed on a BD Accuri™ C6 Flow Cytometer (BD Biosciences) and FlowJo software (RRID:SCR_008520).

### RNA extraction and RT-qPCR

For RNA analysis, total RNA from cells was extracted using either NZYol (NZYTech) or GRS Total RNA Kit (GRiSP), according to manufacturer’s instructions and samples were reverse transcribed using NZY First-Strand cDNA Synthesis (NZYtech), according to the manufacturer’s instructions. Quantitative real-time PCR (qPCR) was performed using NZYSupreme qPCR Green Master Mix (2x), (NZYTech) in an ABI 7500 thermocycler. The 2^-ΔΔCt^ method was used to determine gene expression levels. All primer sequences are listed in Supplementary Table S5.

### Western blotting

Whole-cell extracts were prepared using RIPA buffer (Thermo Scientific) containing protease and phosphatase inhibitors (Thermo Scientific). Cell lysates were then centrifuged at 15,100 g for 10 min at 4°C to remove insoluble cellular components. Histone-enriched (acidic) proteins extracts were prepared using Histone Extraction Kit (Abcam ab113476) following manufacturer instructions. Protein concentration was measured using Pierce BCA Protein Assay Kit (Thermo Scientific) and a plate-reader Tecan Infinite M200 (Tecan). Samples were prepared by boiling with NuPAGE LDS sample buffer (Invitrogen) for 5 min at 95°C in the presence of 50 mM DTT and protein standard (NZYTech, MB09002) was used. 5-15 μg of total protein and 5-7.5 μg for histone-enriched) were subjected to SDS-PAGE in, respectively, homemade 10% and 17% Tris-glycine SDS-PAGE gels. Proteins were wet-transferred onto a nitrocellulose membrane (Amersham), in transfer buffer containing Tris-glycine and methanol (20% v/v). After transfer, the membrane was stained with Ponceau S (0.1% (w/v), Carl Roth) and subsequently blocked with 5% BSA (prepared in TBS-tween20 (0.05%), for 1h at RT. Membranes were incubated overnight at 4°C,with mouse anti-PGC-1α (Calbiochem, KP9803, 1:1000), mouse anti-PINK1 (Santa Cruz Biotechnology, sc-517353, 1:500), rabbit anti-p62 (Cell Signaling 5114, 1:1000), rabbit anti-histone H3 (Abcam ab1791, 1:2000), rabbit anti-pan histone H3 (acetyl K9 + K14 + K18 + K23 + K27) (Abcam ab47915, 1:1000), followed by incubation with either anti-mouse or anti-rabbit HRP-conjugated secondary antibodies (Invitrogen, G21040 and G21234, respectively, both 1:10000). Membranes containing whole-cell lysates were also incubated with an HRP-conjugated rabbit anti-β-actin (Invitrogen MAB-32540, 1:1000). The Pierce ECL (Amersham, RPN2235) detection system and Chemidoc (ChemiDoc Touch Imaging System, Bio-Rad) were used. Quantification of densitometric units was performed using Image Lab Ver. 6.1 software (RRDI:SCR_014210) and uncropped membranes are presented at Figure S5.

### Seahorse XFe96 Extracellular Flux Analysis

Transduced MEFs and AEFs (mock and MGT) on D11, as well as HL-1 cells, were dissociated, counted and plated in the Seahorse XF96 Cell Culture Microplates (Bioscience), coated with 0.1% gelatin, at a density of 10,000 cells/well. On the next day, the Seahorse Cell Mito Stress Test profile was performed using the Seahorse XFe96 Extracellular Flux Analyzer (Agilent Scientific Instruments, California, USA) (Amorim et al., 2022). Briefly, cell culture medium from the plates was replaced by 175 µL/well of pre-warmed low-buffered serum-free minimal DMEM (D5030, SIGMA-Aldrich, USA) medium supplemented with 1 mM pyruvate, 2 mM glutamine, and 10 mM glucose (pH 7.4), and cells were incubated in a non-CO_2_ incubator at 37°C for 1 h. Oxygen consumption rate (OCR) was measured under standard conditions and after the sequential addition of oligomycin (2 μM) (port A), BAM-15 (2 μM) (port B) and rotenone (1 μM) plus antimycin A (1 μM) (port C). Three baseline rate measurements of cell OCR were made using a 3 min mix, 3 min measure cycle. The compounds were then pneumatically injected by the XFe96 Analyzer into each well, mixed and OCR measurements made using a 3 min mix, 3 min measure cycle. Results were normalized by cell mass density through the sulforhodamine B (SRB) assay for cell mass determination. Briefly, cells were fixed with 10% (w/v) TCA overnight at 4°C, and then plates were dried in an oven at 37°C. 100 µL of 0.05% SRB in 1% acetic acid solution was added and incubated at 37°C for 1 h. The wells were then washed with 1% acetic acid in water and dried. Then, 100 µL of Tris (pH 10) was added and optical density was measured at 540 nm in Biotek Cytation 3 reader (Biotek Instruments, Winooski, VT, USA) and analyzed using the Wave Desktop 2.6.1 software (RRID:SCR_014526).

### Immunofluorescence and microscopy imaging

#### Mitochondrial morphology analysis

After direct cardiac conversion, cells were dissociated and plated at 2×10^4^ cells per well in 24-well plates with 0.1% gelatin-coated glass coverslips and fixed with PFA (3.7%, Thermo Scientific) followed by permeabilization with 0.1% Triton X-100 (SIGMA) and blocking with 1% BSA (SIGMA) in PBS 1x for 1h at RT. Primary anti-mouse HSP-60 (BD Biosciences, 1:200), anti-rabbit TOM20 (Proteintech, 1:100), or anti-mouse cardiac Troponin T (Abcam, 1:150) antibodies were incubated at 4°C overnight followed by secondary antibodies anti-rabbit Alexa 488 (Invitrogen, 1:500), anti-mouse Alexa 594 (Invitrogen, 1:500), or α-GFP Alexa 488 (Invitrogen, 1:500) for 1h at RT and cells were mounted using Vectashield with DAPI (vectorlabs). Images were acquired using a 63x oil objective of the Zeiss LSM 880 confocal fluorescence microscope.

The mitochondrial network and morphology were evaluated with ImageJ (RRID:SCR_003070, cell counter plugin) plug-in Mitochondrial Analyzer (Chaudhry et al., 2020). Briefly, a z-stack of images (with an interval of 1,26 μm) was acquired and maximum intensity projections were obtained from each z-stack. For each experiment, the same threshold was applied across groups and analysis was performed blinded. The corrected total cell fluorescence (CTCF) was analyzed in ImageJ (SCR_003070, cell counter plugin) and calculated according to: CTCF = Integrated Density – (Area of selected cell x Mean fluorescence of background readings).

#### MitoTracker red and CellROX staining

After direct cardiac conversion, cells were dissociated, counted and plated at a density of 10.4×10^3^ cells on 0.1% gelatin-coated 8-well ibidi plates (plastic ibidi 81506) incubated at 37°C with 5% CO_2_, until the next day. Probe staining was performed directly on the corresponding plates, with CellROX Deep Red (ThermoFisher C10422, 5 μM) and MitoTracker Red (Invitrogen M7512, 150 nM) in HBSS 1x (Gibco) for 30 min at 37°C in the dark. Cells were fixed with PFA (3.7%, Thermo Scientific), for 20 min, washed and mounted using Vectashield with DAPI (Vectorlabs).

For immunostaining combined with MitoTracker red, cells were fixed with PFA (3.7%, Thermo Scientific), followed by permeabilization with 0.5% Triton X-100 (SIGMA) and blocking with 5% goat serum (Abbkine Scientific) and 0.3% Triton X-100 in PBS1x for 1h at RT with agitation. Primary rabbit anti-LC3 (Cell Signalling #3868, 1:600) antibody was incubated at 4°C overnight followed by secondary antibody anti-rabbit Alexa 488 (ThermoFisher Scientific, 1:600) for 1h at RT. Vectashield mounting medium containing DAPI (ref) was added directly into each well. The images were obtained through a 63x oil objective of the Zeiss LSM 880 confocal fluorescence microscope and analyzed with ImageJ software (SCR_003070). After determining cell boundary, the fluorescence intensities of CellROX, MitoTracker red and LC3 were measured as integrated density (IntDen). Colocalization analyses were performed on fields of view using the Pearson’s coefficient on the JACoP plug-in. Briefly, this coefficient describes how well different fluorochromes are related by a linear equation and goes from −1 (negative correlation) to 1 (positive correlation and perfect colocalization), with 0 standing for no correlation.

#### TMRE live cell imaging

After direct cardiac conversion, cells were dissociated, counted and plated at a density of 10.4×10^3^ cells on 0.1% gelatin-coated 8-well ibidi plates (plastic ibidi 81506) and incubated at 37°C with 5% CO_2_, until the next day.

For negative control, medium was removed and FCCP (Abcam, 20μM) was added for 10 min at 37°C. TMRE Red (Abcam, ab113852, 50 nM), prepared in growth medium, was added for 20min at 37°C in the dark, cells were washed and imaged. Images were captured utilizing a 63x oil objective on the Zeiss LSM 880 confocal fluorescence microscope. The mean fluorescence intensity levels were analysed in the Mitometer software, an algorithm for fast, unbiased, and automated segmentation and tracking of mitochondria in live cells. (Lefebvre et al., 2021) The outputs given by Mitometer were analyzed, under optimized conditions, providing the observed result per field-of-view (Vitória et al., 2023).

### Metabolite extraction and analysis by Liquid Chromatography-Mass Spectrometry (LC–MS)

Transduced MEFs (mock and MGT) cells on day 11, as well as HL-1 cells, about 4×10^6^, were used, in order to extract metabolites and soluble lipids, with 80% (v/v) methanol (at −80°C or on dry ice or liquid nitrogen) as previously described. (M. Yuan et al., 2012) Pellets were kept at −80°C until extraction. The dried samples were resuspended in 50 μl of LC/MS phase A containing an internal standard (Leu-Tyr, 0.02 mg/mL, SIGMA). Quality control (QC) samples, created by pooling aliquots of all samples, were produced. Five microliters of each sample were analyzed by C18-LC-MS using an Ascentis® Express 90 Å C18 HPLC column (15 cm x 2.1 mm; 2.7 μm, Supelco®) with an HPLC system (Ultimate 3000 Dionex, Thermo Fisher Scientific, Bremen, Germany) coupled online to a Q-Exactive™ Mass Spectrometer (Thermo Fisher Scientific, Bremen, Germany). The solvent system consisted of two mobile phases, a mobile phase A (0.1% formic acid, 2% acetonitrile in water) and mobile phase B (B: 0.1% formic acid, 100% acetonitrile. Metabolites were eluted with a 100% of mobile phase A for 2 min, followed by a linear gradient of 0–100% B over 18 min and held for 2 min at a flow rate of 260 mL/min and a temperature of 35°C. The mass spectrometer (MS) was operated as previously described. (Maurício et al., 2022) Positive and negative ion modes were employed (voltages 3.1 kV and −2.8 kV, respectively). The capillary temperature was set to 350 °C, and a sheath gas flow of 20 U was used. MS survey scans were conducted with a resolution of 70,000, an AGC target of 1 × 10^6^, and a maximum IT of 100 ms. The scan range spanned from 65 to 900 m/z (mass-to-charge ratio). From each MS scan, the 10 most intense peaks were selected for HCD MS/MS experiments at a resolution of 17,500, an AGC target of 10^3^, and a maximum IT of 50 ms and a dynamic exclusion time of 30 s. The normalized collision energy was adjusted to 20, 30, and 40, and an isolation width of 1.5 Th (Thomson) was used. The Xcalibur data system (v3.3, Thermo Fisher Scientific, Waltham, MA, USA) was utilized for data acquisition. Metabolites were identified using MS-DIAL v 4.70 software (Mass Spectrometry-Data Independent Analysis (MS-DIAL). (Tsugawa et al., 2020) The identification of metabolites was conducted in both negative and positive ionization modes, utilizing the QC raw files obtained from the MS/MS analysis for each mode. Validated species were integrated and quantified using MZmine v 2.53 software, (Pluskal et al., 2010) and depicted in Supplementary Table S2. To normalize the data, the peak areas of the metabolites and lipid precursor’s extracted ion chromatograms (XIC) were normalized using generalized log2 and autoscaling. Univariate and multivariate statistical analyses were carried out using R version 4.2.3 in Rstudio version 2022.12.0. Principal component analysis (PCA) was conducted using the R libraries FactoMineR, (Lê et al., 2008) and factoextra. Heatmaps were generated using the R package pheatmap (RRID:SCR_016418), with "Euclidean" as clustering distance, and "ward.D" as the clustering method. The R package ggplot2, (Wickham, 2008) (RRID:SCR_014601) was used to create graphics and boxplots. To evaluate the significance differences between conditions, ANOVA and Kruskal-Wallis tests were applied, complimented with a Tukey’s or Dunn’s test, respectively and the data normality was confirmed using the Shapiro-Wilk test. All tests were done resorting to the R package Rstatix, using a p-value below 0.05 to consider statistically significant. Metaboanlyst was used to perform metabolic pathway enrichment analysis and pathway topology analysis.

### Histone extraction and analysis of acetylation and methylation marks by LC–MS

Histone were enriched from 1-2×10^6^ transduced MEFs and AEFs (mock and MGT) cells on day 11, adult mouse cardiomyocyte cell line HL-1 or primary neonatal mouse cardiac fibroblasts (NMCF) or cardiomyocytes (NMCM) using the Histone extraction kit (Abcam) according to the manufacturer’s instructions. Protein extracts were resuspended in Balanced-DTT Buffer and stored at −80°C until analysis. Histone protein concentration and purity was accessed by BCA protein quantification assay (Thermo Scientific) and SDS-PAGE analysis with 17% polyacrylamide gel and Coomassie staining (0.12% (w/v) Coomassie G-250 in 20% methanol (v/v)) using known amounts of histone H3.1 recombinant protein (EpiCypher, Inc). Approximately 4 µg of histone octamer were mixed with an equal amount of heavy-isotope labelled histones, which were used as an internal standard, (R Noberini & Bonaldi, 2017) and separated on a 17% SDS-PAGE gel. Histone bands were excised, chemically acylated with propionic anhydride and in-gel digested with trypsin, followed by peptide N-terminal derivatization with phenyl isocyanate (PIC). (Roberta Noberini et al., 2021) Peptide mixtures were separated by reversed-phase chromatography on an EASY-Spray column (Thermo Fisher Scientific), 25-cm long (inner diameter 75 µm, PepMap C18, 2 µm particles), which was connected online to a Q Exactive Plus instrument (Thermo Fisher Scientific) through an EASY-Spray™ Ion Source (Thermo Fisher Scientific), as described (2). The acquired RAW data were analyzed using EpiProfile 2.0, (Z.-F. Yuan et al., 2018) followed by manual validation. For each histone modified peptide, a % relative abundance (%RA) value for the sample (light channel - L) or the internal standard (heavy channel - H) was estimated by dividing the area under the curve of each acetylated peptide for the sum of the areas corresponding to all the observed forms of that peptide and multiplying by 100. Light/Heavy (L/H) ratios of %RA were then calculated and are reported in Supplementary Table S1 and S4. Data analysis and visualization were performed using Perseus (RRID:SCR_015753) and GraphPad Prism (RRID:SCR_002798). The mass spectrometry data have been deposited to the ProteomeXchange Consortium, (Vizcaíno et al., 2014) via the PRIDE partner repository with the dataset identifier PXD046542.

### Quantificaton and statistical analysis

All experiments were carried out in three or more biological replicates. Quantifications of data and statistical analysis of metabolomics and histone proteomics data are described under various Method details sections where applicable. Graphical data denote the mean ± s.d or mean ± s.e.m (of n = 3 or more biological replicates, cells quantified or mice) and are depicted by column graph scatter dot plot using GraphPad Prism 8.4.0 software (SCR_002798) or as otherwise specified. Details of the sample numbers can be found in the appropriate figure legend. Unless indicated otherwise, the P values for analyses was determined by two-tailed Student’s *t*-test between two groups, and one-way or two-way analysis of variance (ANOVA) followed by Tukey’s post-test for multiple comparisons or Kruskal Wallis with a false discovery rate correction (with an FDR value of 0.05) using a GraphPad Prism 8.4.0 software. For normality, the Shapiro-Wilk test was performed.∗, p<0.05; ∗∗, p<0.01; ∗∗∗, p<0.001; ∗∗∗*, p<0.0001.

## AUTHOR CONTRIBUTIONS

F.S developed experimental approaches and methodology, performed majority of experiments and data analysis. M.C., R.D. and B.B. performed experiments and data analysis. R.S.F. established methodology in mitochondria network analysis. R.N. and T.B. performed MS-based histone proteomics. P.D performed LC-MS based metabolomics. J.T., P.O., C.B., and B.B. de J. performed experiments, provided tools, or analyzed data. P.D, B.B. de J. and S.N.-P. acquired funding; S.N.-P. conceptualized and administered the project and wrote the manuscript with help from all authors.

## ACKNOWLEDGMENTS

The authors thank Dr. Julia Leonardy for performing NMCF and NMCM isolation; Tatiana Maurício and Chiara Ghirardi for technical assistance; Dr. Rui Vitorino for guided assistance in histone proteomics analysis; Dr. Rita Ferreira, Dr. Bruno Neves, Dr. Catarina Almeida, Dr. Luisa Helguero and Dr. Margarida Fardilha for providing reagents; Dr. Diogo Trigo and José J. M. Vitória for assistance in the analysis on Mitometer. Other support: iBiMED Cell Culture and Light Microscopy Facility (a node of PPBI, Portuguese Platform of BioImaging, POCI-01-0145-FEDER-022122) and iBiMED animal facility S. N.-P. and B.B. de J. reports funds from Fundação para a Ciência e Tecnologia (FCT EXPL/BIA-CEL/0358/2021, 2022.01199.PTDC DOI 10.54499/2022.01199.PTDC), Portugal, during the conduct of the study. The MS-based histone modification analyses were performed thanks to the support of EPIC-XS, project number 823839, funded by the Horizon 2020 programme of the European Union. The LC-MS based metabolomic studies were performed at Aveiro University Mass Spectrometry Center. S. N.-P. received funding from FCT CEECIND Program (2020.00355.CEECIND) and J. T. acknowledges FCT, I.P. for the research contract 2020.01560.CEECIND. F. S. and M.C. are recipients of FCT predoctoral fellowship (SFRH/BD/146204/2019 and UI/BD/151373/2021, respectively), Portugal. No disclosures were reported by the other authors.

## DATA AVAILABILITY STATEMENT

LC-MS based metabolomics and histone proteomics data from this work and uncropped immunoblots are available in supplemental information (Table S1-S4 and Supplementary Figure S4 and S7). The LC-MS based histone proteomics data have been deposited in the ProteomeXchange Consortium PRIDE partner repository with the dataset identifier PXD046542. This paper does not report original code. Any additional information required to reanalyze the data reported in this paper is available from the lead contact upon request.

## REFERENCES

Amorim, R., Cagide, F., Tavares, L. C., Simões, R. F., Soares, P., Benfeito, S., Baldeiras, I., Jones, J. G., Borges, F., Oliveira, P. J., & Teixeira, J. (2022). Mitochondriotropic antioxidant based on caffeic acid AntiOxCIN(4) activates Nrf2-dependent antioxidant defenses and quality control mechanisms to antagonize oxidative stress-induced cell damage. Free Radical Biology & Medicine, 179, 119–132. 10.1016/j.freeradbiomed.2021.12.304

Bernardes de Jesus, B., Marinho, S. P., Barros, S., Sousa-Franco, A., Alves-Vale, C., Carvalho, T., & Carmo-Fonseca, M. (2018). Silencing of the lncRNA Zeb2-NAT facilitates reprogramming of aged fibroblasts and safeguards stem cell pluripotency. Nature Communications, 9(1), 94. 10.1038/s41467-017-01921-6

Cardoso, A. C., Lam, N. T., Savla, J. J., Nakada, Y., Pereira, A. H. M., Elnwasany, A., Menendez-Montes, I., Ensley, E. L., Bezan Petric, U., Sharma, G., Sherry, A. D., Malloy, C. R., Khemtong, C., Kinter, M. T., Tan, W. L. W., Anene-Nzelu, C. G., Foo, R. S. Y., Nguyen, N. U. N., Li, S., … Sadek, H. A. (2020). Mitochondrial substrate utilization regulates cardiomyocyte cell-cycle progression. Nature Metabolism, 2(2), 167–178. 10.1038/s42255-020-0169-x

Carey, B. W., Finley, L. W. S., Cross, J. R., Allis, C. D., & Thompson, C. B. (2015). Intracellular α-ketoglutarate maintains the pluripotency of embryonic stem cells. Nature, 518(7539), 413–416. 10.1038/nature13981

Carrer, A., Parris, J. L. D., Trefely, S., Henry, R. A., Montgomery, D. C., Torres, A., Viola, J. M., Kuo, Y.-M., Blair, I. A., Meier, J. L., Andrews, A. J., Snyder, N. W., & Wellen, K. E. (2017). Impact of a High-fat Diet on Tissue Acyl-CoA and Histone Acetylation Levels. The Journal of Biological Chemistry, 292(8), 3312–3322. 10.1074/jbc.M116.750620

Chaudhry, A., Shi, R., & Luciani, D. S. (2020). A pipeline for multidimensional confocal analysis of mitochondrial morphology, function, and dynamics in pancreatic β-cells. American Journal of Physiology. Endocrinology and Metabolism, 318(2), E87–E101. 10.1152/ajpendo.00457.2019

Claycomb, W. C., Lanson, N. A. J., Stallworth, B. S., Egeland, D. B., Delcarpio, J. B., Bahinski, A., & Izzo, N. J. J. (1998). HL-1 cells: a cardiac muscle cell line that contracts and retains phenotypic characteristics of the adult cardiomyocyte. Proceedings of the National Academy of Sciences of the United States of America, 95(6), 2979–2984. 10.1073/pnas.95.6.2979

Cornacchia, D., Zhang, C., Zimmer, B., Chung, S. Y., Fan, Y., Soliman, M. A., Tchieu, J., Chambers, S. M., Shah, H., Paull, D., Konrad, C., Vincendeau, M., Noggle, S. A., Manfredi, G., Finley, L. W. S., Cross, J. R., Betel, D., & Studer, L. (2019). Lipid Deprivation Induces a Stable, Naive-to-Primed Intermediate State of Pluripotency in Human PSCs. Cell Stem Cell, 25(1), 120–136.e10. 10.1016/j.stem.2019.05.001

Costa, A., Cushman, S., Haubner, B. J., Derda, A. A., Thum, T., & Bär, C. (2022). Neonatal injury models: integral tools to decipher the molecular basis of cardiac regeneration. Basic Research in Cardiology, 117(1), 26. 10.1007/s00395-022-00931-w

Ferrari, K. J., Scelfo, A., Jammula, S., Cuomo, A., Barozzi, I., Stützer, A., Fischle, W., Bonaldi, T., & Pasini, D. (2014). Polycomb-dependent H3K27me1 and H3K27me2 regulate active transcription and enhancer fidelity. Molecular Cell, 53(1), 49–62. 10.1016/j.molcel.2013.10.030

Fu, Y., Huang, C., Xu, X., Gu, H., Ye, Y., Jiang, C., Qiu, Z., & Xie, X. (2015). Direct reprogramming of mouse fibroblasts into cardiomyocytes with chemical cocktails. Cell Research, 25(9), 1013–1024. 10.1038/cr.2015.99

Garry, G. A., Bezprozvannaya, S., Chen, K., Zhou, H., Hashimoto, H., Morales, M. G., Liu, N., Bassel-Duby, R., & Olson, E. N. (2021). The histone reader PHF7 cooperates with the SWI/SNF complex at cardiac super enhancers to promote direct reprogramming. Nature Cell Biology, 23(5), 467–475. 10.1038/s41556-021-00668-z

Gong, G., Song, M., Csordas, G., Kelly, D. P., Matkovich, S. J., & Dorn, G. W. 2nd. (2015). Parkin-mediated mitophagy directs perinatal cardiac metabolic maturation in mice. Science (New York, N.Y.), 350(6265), aad2459. 10.1126/science.aad2459

Hashimoto, H., Olson, E. N., & Bassel-Duby, R. (2018). Therapeutic approaches for cardiac regeneration and repair. Nature Reviews Cardiology, 15(10), 585–600. 10.1038/s41569-018-0036-6

Hirai, H., & Kikyo, N. (2014). Inhibitors of suppressive histone modification promote direct reprogramming of fibroblasts to cardiomyocyte-like cells. Cardiovascular Research, 102(1), 189–190. 10.1093/cvr/cvu023

Hong, X., Isern, J., Campanario, S., Perdiguero, E., Ramírez-Pardo, I., Segalés, J., Hernansanz-Agustín, P., Curtabbi, A., Deryagin, O., Pollán, A., González-Reyes, J. A., Villalba, J. M., Sandri, M., Serrano, A. L., Enríquez, J. A., & Muñoz-Cánoves, P. (2022). Mitochondrial dynamics maintain muscle stem cell regenerative competence throughout adult life by regulating metabolism and mitophagy. Cell Stem Cell, 29(9), 1298–1314.e10. 10.1016/j.stem.2022.07.009

Huang, C., Tu, W., Fu, Y., Wang, J., & Xie, X. (2018). Chemical-induced cardiac reprogramming in vivo. Cell Research, 28(6), 686–689. 10.1038/s41422-018-0036-4

Ieda, M., Fu, J., Delgado-olguin, P., Vedantham, V., Hayashi, Y., Bruneau, B. G., & Srivastava, D. (2010). Direct Reprogramming of Fibroblasts into Functional Cardiomyocytes by Defined Factors. Cell, 142(3), 375–386. 10.1016/j.cell.2010.07.002

Ifkovits, J. L., Addis, R. C., Epstein, J. A., & Gearhart, J. D. (2014). Inhibition of TGFβ Signaling Increases Direct Conversion of Fibroblasts to Induced Cardiomyocytes. PLOS ONE, 9(2), 1–11. 10.1371/journal.pone.0089678

Intlekofer, A. M., & Finley, L. W. S. (2019). Metabolic signatures of cancer cells and stem cells. Nature Metabolism, 1(2), 177–188. 10.1038/s42255-019-0032-0

Lazarou, M., Sliter, D. A., Kane, L. A., Sarraf, S. A., Wang, C., Burman, J. L., Sideris, D. P., Fogel, A. I., & Youle, R. J. (2015). The ubiquitin kinase PINK1 recruits autophagy receptors to induce mitophagy. Nature, 524(7565), 309–314. 10.1038/nature14893

Lê, S., Josse, J., & Husson, F. (2008). FactoMineR: An R Package for Multivariate Analysis. Journal of Statistical Software, 25(1), 1–18. 10.18637/jss.v025.i01

Lefebvre, A. E. Y. T., Ma, D., Kessenbrock, K., Lawson, D. A., & Digman, M. A. (2021). Automated segmentation and tracking of mitochondria in live-cell time-lapse images. Nature Methods, 18(9), 1091–1102. 10.1038/s41592-021-01234-z

Lehman, J. J., & Kelly, D. P. (2002). Transcriptional activation of energy metabolic switches in the developing and hypertrophied heart. Clinical and Experimental Pharmacology & Physiology, 29(4), 339–345. 10.1046/j.1440-1681.2002.03655.x

Liu, Z., Chen, O., Zheng, M., Wang, L., Zhou, Y., Yin, C., Liu, J., & Qian, L. (2016). Re-patterning of H3K27me3, H3K4me3 and DNA methylation during fibroblast conversion into induced cardiomyocytes. Stem Cell Research, 16(2), 507–518. 10.1016/j.scr.2016.02.037

Liu, Z., Wang, L., Welch, J. D., Ma, H., Zhou, Y., Vaseghi, H. R., Yu, S., Wall, J. B., Alimohamadi, S., Zheng, M., Yin, C., Shen, W., Prins, J. F., Liu, J., & Qian, L. (2017). Single-cell transcriptomics reconstructs fate conversion from fibroblast to cardiomyocyte. Nature, 551(7678), 100–104. 10.1038/nature24454

Lopaschuk, G. D., Collins-Nakai, R. L., & Itoi, T. (1992). Developmental changes in energy substrate use by the heart. Cardiovascular Research, 26(12), 1172–1180. 10.1093/cvr/26.12.1172

Lu, D., Chatterjee, S., Xiao, K., Riedel, I., Huang, C.-K., Costa, A., Cushman, S., Neufeldt, D., Rode, L., Schmidt, A., Juchem, M., Leonardy, J., Büchler, G., Blume, J., Gern, O.-L., Kalinke, U., Wen Tan, W. L., Foo, R., Vink, A., … Thum, T. (2022). A circular RNA derived from the insulin receptor locus protects against doxorubicin-induced cardiotoxicity. European Heart Journal, 43(42), 4496–4511. 10.1093/eurheartj/ehac337

Lynch, C. J., Bernad, R., Martínez-Val, A., Shahbazi, M. N., Nóbrega-Pereira, S., Calvo, I., Blanco-Aparicio, C., Tarantino, C., Garreta, E., Richart-Ginés, L., Alcazar, N., Graña-Castro, O., Gómez-Lopez, G., Aksoy, I., Muñoz-Martín, M., Martinez, S., Ortega, S., Prieto, S., Simboeck, E., … Serrano, M. (2020). Global hyperactivation of enhancers stabilizes human and mouse naive pluripotency through inhibition of CDK8/19 Mediator kinases. Nature Cell Biology. 10.1038/s41556-020-0573-1

Maryanovich, M., & Gross, A. (2013). A ROS rheostat for cell fate regulation. Trends in Cell Biology, 23(3), 129–134. 10.1016/j.tcb.2012.09.007

Maurício, T., Aveiro, S., Guedes, S., Lopes, D., Melo, T., Neves, B. M., Domingues, R., & Domingues, P. (2022). Multi-Omic Profiling of Macrophages Treated with Phospholipids Containing Omega-3 and Omega-6 Fatty Acids Reveals Complex Immunomodulatory Adaptations at Protein, Lipid and Metabolic Levels. International Journal of Molecular Sciences, 23(4). 10.3390/ijms23042139

McNally, L. A., Altamimi, T. R., Fulghum, K., & Hill, B. G. (2021). Considerations for using isolated cell systems to understand cardiac metabolism and biology. Journal of Molecular and Cellular Cardiology, 153(July 2020), 26–41. 10.1016/j.yjmcc.2020.12.007

Menendez-Montes, I., Abdisalaam, S., Xiao, F., Lam, N. T., Mukherjee, S., Szweda, L. I., Asaithamby, A., & Sadek, H. A. (2021). Mitochondrial fatty acid utilization increases chromatin oxidative stress in cardiomyocytes. Proceedings of the National Academy of Sciences of the United States of America, 118(34), 3–5. 10.1073/pnas.2101674118

Noberini, R, & Bonaldi, T. (2017). A Super-SILAC Strategy for the Accurate and Multiplexed Profiling of Histone Posttranslational Modifications. Methods in Enzymology, 586, 311–332. 10.1016/bs.mie.2016.09.036

Noberini, Roberta, Savoia, E. O., Brandini, S., Greco, F., Marra, F., Bertalot, G., Pruneri, G., McDonnell, L. A., & Bonaldi, T. (2021). Spatial epi-proteomics enabled by histone post-translational modification analysis from low-abundance clinical samples. Clinical Epigenetics, 13(1), 145. 10.1186/s13148-021-01120-7

Nóbrega-Pereira, S., Caiado, F., Carvalho, T., Matias, I., Graça, G., Gonçalves, L. G., Silva-Santos, B., Norell, H., & Dias, S. (2018). VEGFR2-Mediated Reprogramming of Mitochondrial Metabolism Regulates the Sensitivity of Acute Myeloid Leukemia to Chemotherapy. Cancer Research, 78(3), 731– 741. 10.1158/0008-5472.CAN-17-1166

Nóbrega-Pereira, S., & de Jesus, B. B. (2020). Cellular Reprogramming and Aging. 73–91. 10.1007/978-3-030-43939-2_5

Pal, S., & Tyler, J. K. (2016). Epigenetics and aging. Science Advances, 2(7), e1600584. 10.1126/sciadv.1600584

Paquin, K. L., & Howlett, N. G. (2018). Understanding the Histone DNA Repair Code: H4K20me2 Makes Its Mark. Molecular Cancer Research, 16(9), 1335–1345. 10.1158/1541-7786.MCR-17-0688

Pascual, G., Domínguez, D., Elosúa-Bayes, M., Beckedorff, F., Laudanna, C., Bigas, C., Douillet, D., Greco, C., Symeonidi, A., Hernández, I., Gil, S. R., Prats, N., Bescós, C., Shiekhattar, R., Amit, M., Heyn, H., Shilatifard, A., & Benitah, S. A. (2021). Dietary palmitic acid promotes a prometastatic memory via Schwann cells. Nature, 599(7885), 485–490. 10.1038/s41586-021-04075-0

Persad, K. L., & Lopaschuk, G. D. (2022). Energy Metabolism on Mitochondrial Maturation and Its Effects on Cardiomyocyte Cell Fate. Frontiers in Cell and Developmental Biology, 10. 10.3389/fcell.2022.886393

Pluskal, T., Castillo, S., Villar-Briones, A., & Orešič, M. (2010). MZmine 2: Modular framework for processing, visualizing, and analyzing mass spectrometry-based molecular profile data. BMC Bioinformatics, 11(1), 395. 10.1186/1471-2105-11-395

Qian, L., Huang, Y., Spencer, C. I., Foley, A., Vedantham, V., Liu, L., Conway, S. J., Fu, J. D., & Srivastava, D. (2012). In vivo reprogramming of murine cardiac fibroblasts into induced cardiomyocytes. Nature, 485(7400), 593–598. 10.1038/nature11044

Ryu, D., Mouchiroud, L., Andreux, P. A., Katsyuba, E., Moullan, N., Nicolet-Dit-Félix, A. A., Williams, E. G., Jha, P., Lo Sasso, G., Huzard, D., Aebischer, P., Sandi, C., Rinsch, C., & Auwerx, J. (2016). Urolithin A induces mitophagy and prolongs lifespan in C. elegans and increases muscle function in rodents. Nature Medicine, 22(8), 879–888. 10.1038/nm.4132

Santos, F., Correia, M., Nóbrega-Pereira, S., & Bernardes de Jesus, B. (2020). Age-Related Pathways in Cardiac Regeneration: A Role for lncRNAs? Frontiers in Physiology, 11, 583191. 10.3389/fphys.2020.583191

Singh, V. P., Pinnamaneni, J. P., Pugazenthi, A., Sanagasetti, D., Mathison, M., Wang, K., Yang, J., & Rosengart, T. K. (2020). Enhanced generation of induced cardiomyocytes using a small-molecule cocktail to overcome barriers to cardiac cellular reprogramming. Journal of the American Heart Association, 9(12). 10.1161/JAHA.119.015686

Song, K., Nam, Y. J., Luo, X., Qi, X., Tan, W., Huang, G. N., Acharya, A., Smith, C. L., Tallquist, M. D., Neilson, E. G., Hill, J. A., Bassel-Duby, R., & Olson, E. N. (2012). Heart repair by reprogramming non-myocytes with cardiac transcription factors. Nature, 485(7400), 599–604. 10.1038/nature11139

Soonpaa, M. H., Kim, K. K., Pajak, L., Franklin, M., & Field, L. J. (1996). Cardiomyocyte DNA synthesis and binucleation during murine development. The American Journal of Physiology, 271(5 Pt 2), H2183-9. 10.1152/ajpheart.1996.271.5.H2183

Tani, H., Sadahiro, T., & Ieda, M. (2018). Direct cardiac reprogramming: A novel approach for heart regeneration. International Journal of Molecular Sciences, 19(9), 1–13. 10.3390/ijms19092629

Tsugawa, H., Ikeda, K., Takahashi, M., Satoh, A., Mori, Y., Uchino, H., Okahashi, N., Yamada, Y., Tada, I., Bonini, P., Higashi, Y., Okazaki, Y., Zhou, Z., Zhu, Z.-J., Koelmel, J., Cajka, T., Fiehn, O., Saito, K., Arita, M., & Arita, M. (2020). A lipidome atlas in MS-DIAL 4. Nature Biotechnology, 38(10), 1159–1163. 10.1038/s41587-020-0531-2

van Gastel, N., Stegen, S., Eelen, G., Schoors, S., Carlier, A., Daniëls, V. W., Baryawno, N., Przybylski, D., Depypere, M., Stiers, P. J., Lambrechts, D., Van Looveren, R., Torrekens, S., Sharda, A., Agostinis, P., Lambrechts, D., Maes, F., Swinnen, J. V., Geris, L., … Carmeliet, G. (2020). Lipid availability determines fate of skeletal progenitor cells via SOX9. Nature, 579(7797), 111–117. 10.1038/s41586-020-2050-1

Vaseghi, H. R., Yin, C., Zhou, Y., Wang, L., Liu, J., & Qian, L. (2016). Generation of an inducible fibroblast cell line for studying direct cardiac reprogramming. Genesis, 54(7), 398–406. 10.1002/dvg.22947

Vitória, J. J. M., de Paula, V., da Cruz E Silva, O. A. B., & Trigo, D. (2023). Optimized Automated Analysis of Live Neuronal Mitochondria Homeostasis Modulation by Isoform-Specific Retinoic Acid Receptors. Journal of Visualized Experiments : JoVE, 197. 10.3791/65452

Vizcaíno, J. A., Deutsch, E. W., Wang, R., Csordas, A., Reisinger, F., Ríos, D., Dianes, J. A., Sun, Z., Farrah, T., Bandeira, N., Binz, P.-A., Xenarios, I., Eisenacher, M., Mayer, G., Gatto, L., Campos, A., Chalkley, R. J., Kraus, H.-J., Albar, J. P., … Hermjakob, H. (2014). ProteomeXchange provides globally coordinated proteomics data submission and dissemination. In Nature biotechnology (Vol. 32, Issue 3, pp. 223–226). 10.1038/nbt.2839

Wang, H., Yang, Y., Liu, J., & Qian, L. (2021). Direct cell reprogramming: approaches, mechanisms and progress. Nature Reviews Molecular Cell Biology, 22(6), 410–424. 10.1038/s41580-021-00335-z

Wang, L., Liu, Z., Yin, C., Asfour, H., Chen, O., Li, Y., Bursac, N., Liu, J., & Qian, L. (2015). Stoichiometry of Gata4, Mef2c, and Tbx5 influences the efficiency and quality of induced cardiac myocyte reprogramming. Circulation Research, 116(2), 237–244. 10.1161/CIRCRESAHA.116.305547

Wang, L., Ma, H., Huang, P., Xie, Y., Near, D., Wang, H., Xu, J., Yang, Y., Xu, Y., Garbutt, T., Zhou, Y., Liu, Z., Yin, C., Bressan, M., Taylor, J. M., Liu, J., & Qian, L. (2020). Down-regulation of Beclin1 promotes direct cardiac reprogramming. Science Translational Medicine, 12(566), 1–23. 10.1126/scitranslmed.aay7856

Wickham, H. (2008). Elegant Graphics for Data Analysis: ggplot2. In Applied Spatial Data Analysis with R.

Yuan, M., Breitkopf, S. B., Yang, X., & Asara, J. M. (2012). A positive/negative ion–switching, targeted mass spectrometry–based metabolomics platform for bodily fluids, cells, and fresh and fixed tissue. Nature Protocols, 7(5), 872–881. 10.1038/nprot.2012.024

Yuan, Z.-F., Sidoli, S., Marchione, D. M., Simithy, J., Janssen, K. A., Szurgot, M. R., & Garcia, B. A. (2018). EpiProfile 2.0: A Computational Platform for Processing Epi-Proteomics Mass Spectrometry Data. Journal of Proteome Research, 17(7), 2533–2541. 10.1021/acs.jproteome.8b00133

Yucel, N., Wang, Y. X., Mai, T., Porpiglia, E., Lund, P. J., Markov, G., Garcia, B. A., Bendall, S. C., Angelo, M., & Blau, H. M. (2019). Glucose Metabolism Drives Histone Acetylation Landscape Transitions that Dictate Muscle Stem Cell Function. Cell Reports, 27(13), 3939–3955.e6. 10.1016/j.celrep.2019.05.092

