## Supplementary figures and images for "Metabolic and age-associated epigenetic barriers during direct reprogramming of mouse fibroblasts into induced cardiomyocytes"

### Supplementary Figure 1

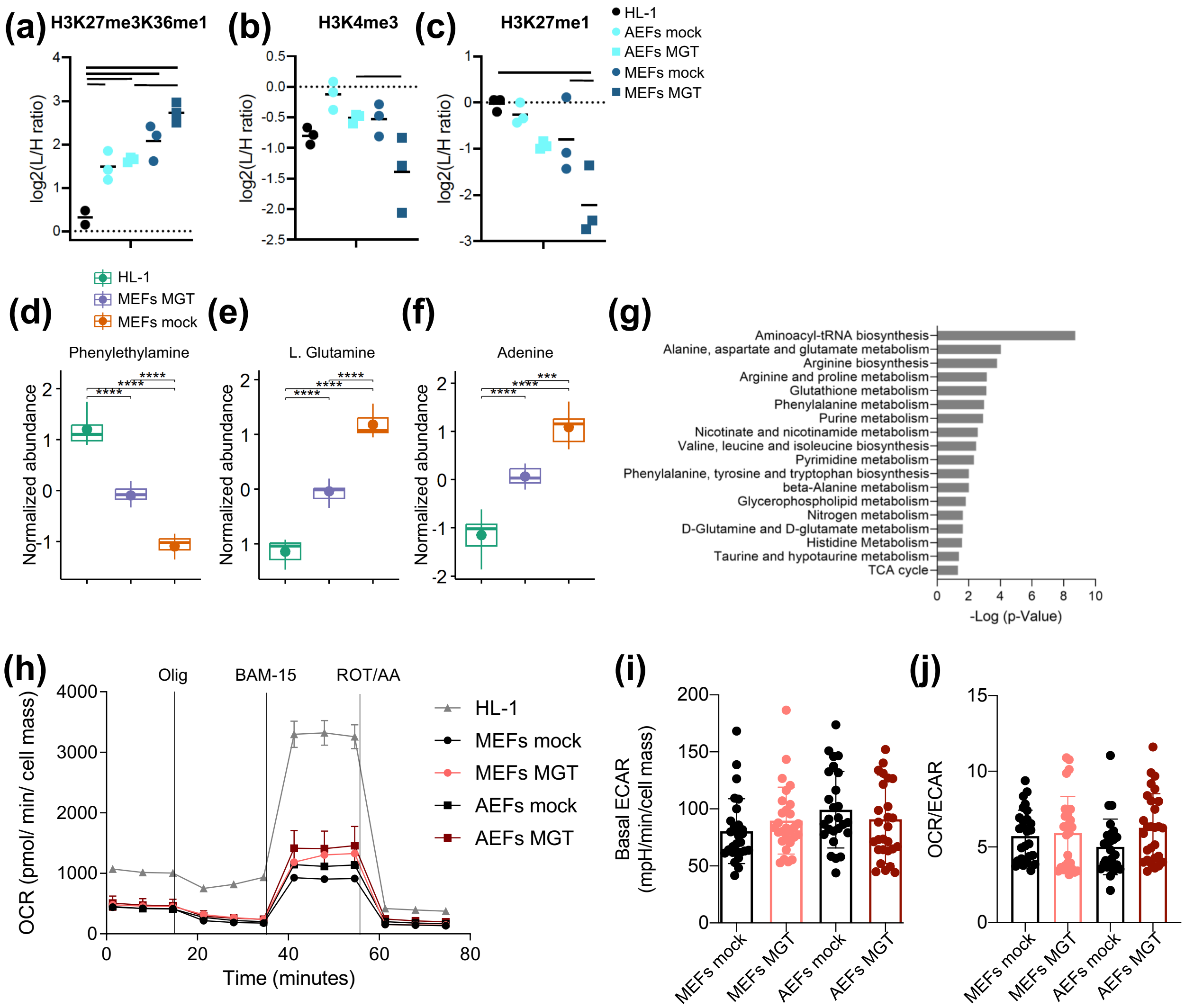

### Supplementary Figure 2

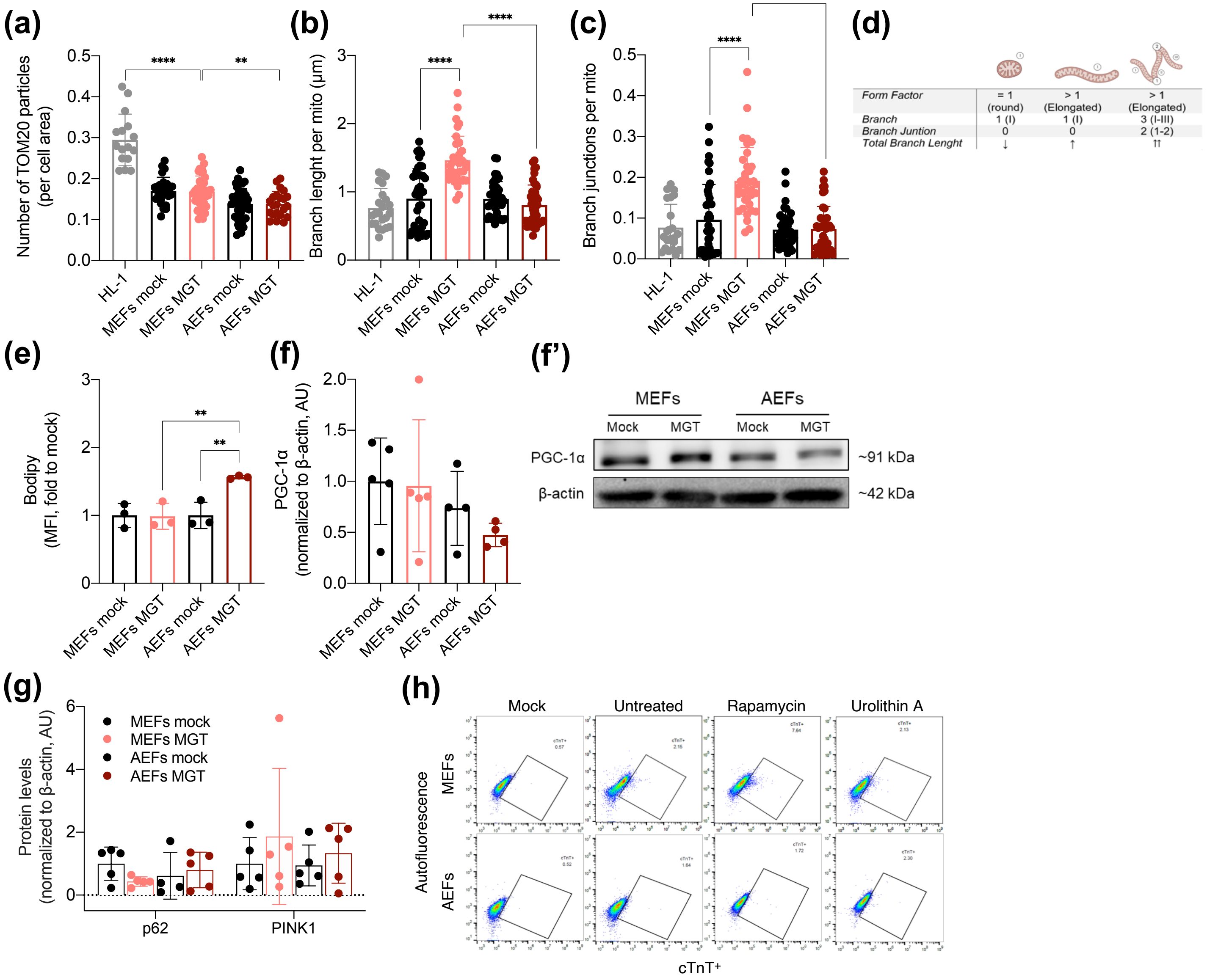

### Supplementary Figure 3

Santos *et al.*, FIGURE S3

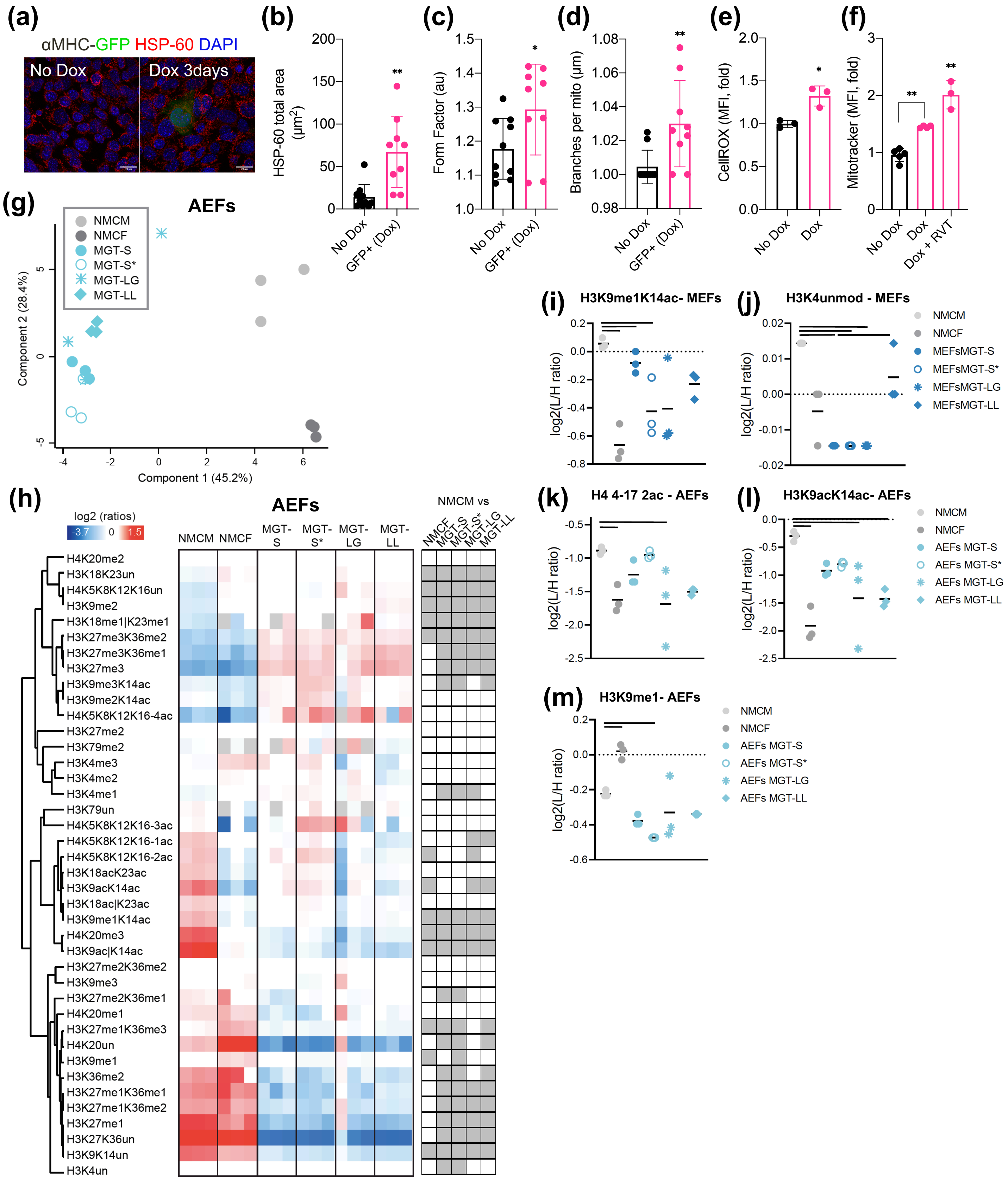

### Supplementary Figure 4

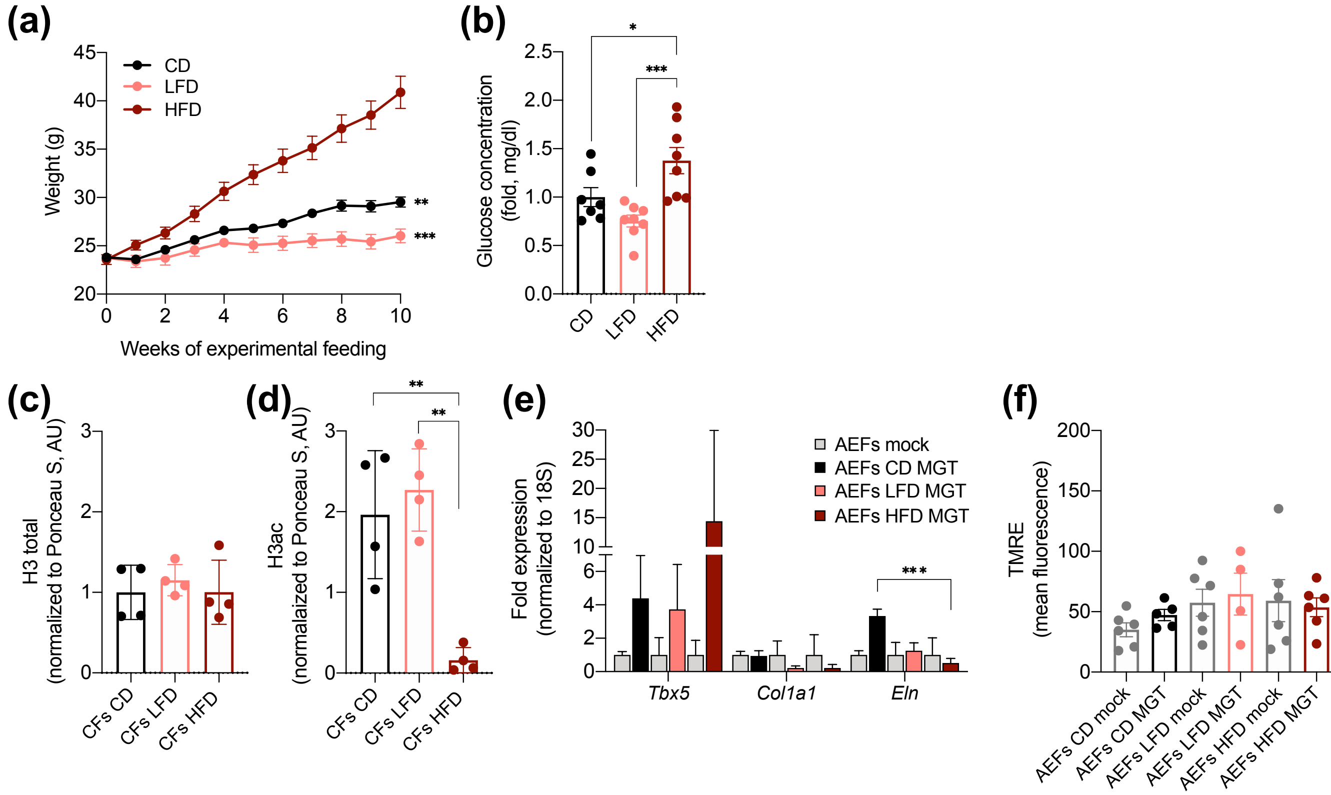
