## Supplementary Figure 5 for "Metabolic and age-associated epigenetic barriers during direct reprogramming of mouse fibroblasts into induced cardiomyocytes"

Santos *et al.*, FIGURE S5

**PGC-1α**

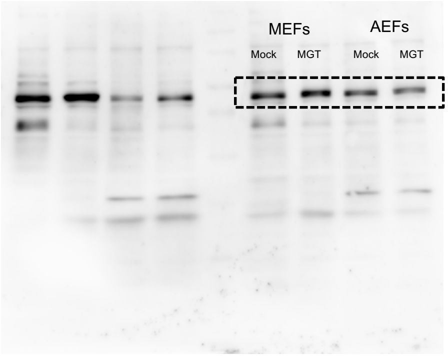


**p62**

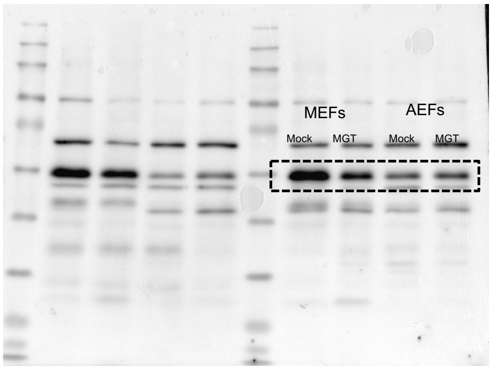


**β-actin**

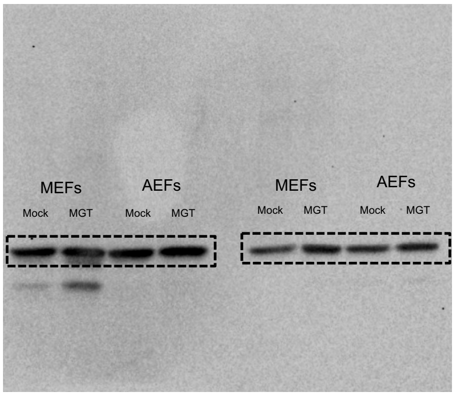


**PINK1**

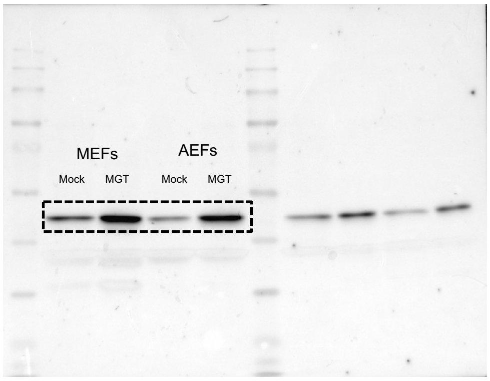


**β-actin**

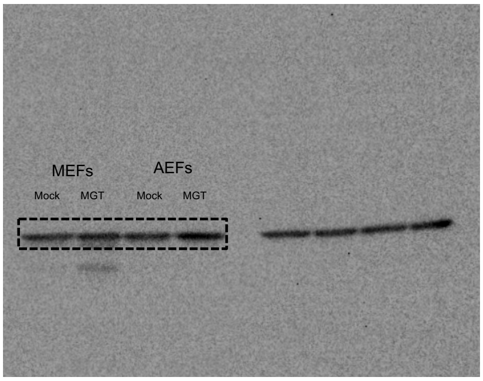


**Ponceau S (Histone H3)**

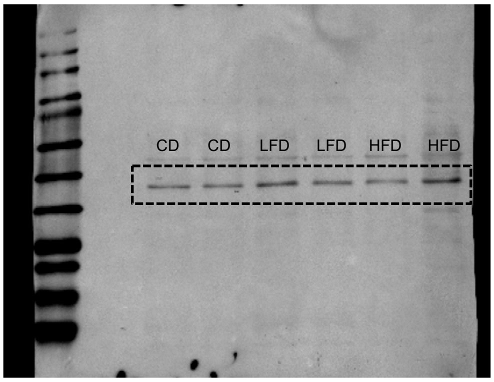


**Ponceau S (H3ac pan-acetyl) pan)**

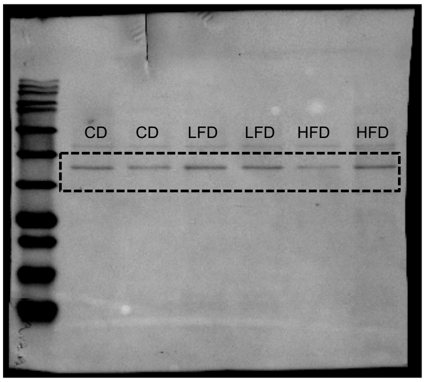


**Histone H3**

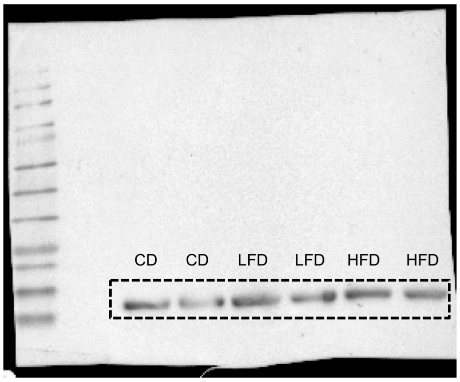


**H3ac pan-acetyl**

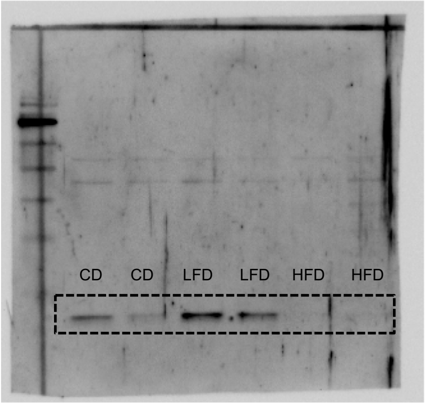
