## Supplementary material for "Metabolic and age-associated epigenetic barriers during direct reprogramming of mouse fibroblasts into induced cardiomyocytes"

**SUPPLEMENTARY FIGURE LEGENDS**

FIGURE S1. Transdifferentiation of mouse fibroblasts into iCMs is accompanied by epigenetic and metabolic transitions. (a-c) Mass spectrometry-based histone acetylation and methylation proteomic bulk analysis in mock and MGT-transduced MEFs and AEFs at D11, and HL-1 mouse cardiomyocyte cell line. Display of selected histone modified peptides showing the levels of tri-methylated form of H3 lysine 27 and mono-methylated lysine 36 (a), tri-methylated form of H3 lysine 4 (b) and mono-methylated form of H3 lysine 27 (c). (d-g) LC-MS-based untargeted metabolomics bulk analysis of MEFs mock or MGT-transduced at D11, and the HL-1 mouse cardiomyocyte cell line. Selected box-plot analysis relative abundances of identified metabolites phenylethylamine (d), L-glutamine (e) and adenine (f) comparison between HL-1 (green boxplots), MEFs mock (orange boxplots) and MEFs MGT (purple boxplots). Pathway analysis of MEFs mock and MGT at D11 displaying bar graph representing the top 18 significantly altered metabolic pathways, in terms of -log(p value) by FDR (g). (h-j) Oxygen consumption rate (OCR) measurement in MEFs and AEFs mock or MGT retroviral-transduced at D11 using Seahorse XF96 Cell Mito Stress Test. OCR measurement (h), basal extracellular acidification rate (ECAR) (i) and OCR/ECAR ratio (j) in MEFs and AEFs mock or MGT retroviral-transduced at D11. Data were normalized to cell mass using the sulforhodamine B (SRB) assay. Each point on the plot indicates individual measurements and mean of n = 3 biological replicates from one representative experiment (a-c), mean ± s.d. of n=6 biological replicates from one representative experiment (d-g) or n=26-28 technical replicates from three independent experiments (h-j).The bar indicates a p-value<0.05 (a-c), each point on the plots (i, j) indicates individual measurements. p values were calculated by one-way ANOVA with False Discovery Rate followed by Tukey’s post-test for multiple comparisons. *p < 0.05; **p < 0.01; ***p < 0.001; ****p < 0.0001 with respect to the indicated groups. Data analysis and visualization were performed using GraphPad Prism (RRID:SCR_002798), Perseus (RRID:SCR_015753) (a-c) and box plots (d-f) with R package ggplot2 (RRID:SCR_014601).

FIGURE S2. Extensive remodeling of mitochondrial network and mitophagy takes place during direct cardiac conversion into iCMs. (a-d) Analysis of number of TOM20 particles (a), branches length (b) and junctions (c) per mitochondria (network connectivity) in confocal microscopy images from MEFs and AEFs mock or MGT-transduced at D11/12 or HL-1 cells. Schematics of mitochondrial morphology parameters quantified using FIJI “*Mitochondrial analyzer*” (d). Morphological and network parameters used to describe mitochondria on a round mitochondrion (left), a long mitochondrion (middle) and mitochondria with multiple branches (I-III) and junctions (1-2) (right). Adapted from (Chaudhry et al., 2020)**.** (e) Flow cytometry quantification of BODIPY 493/503 staining depicted as relative median fluorescence intensity (MFI, fold to corresponding mock) from MEFs and AEFs mock or MGT-transduced at D11/12. (f-f’) Immunoblotting analysis for PGC-1α and β-actin in whole cell extracts densitometric quantification (f, normalized to β-actin) and representative images (f’, uncropped images of blots are shown in Figure S5) from MEFs and AEFs mock or MGT retroviral-transduced at D12. (g) Densitometric quantification of immunoblotting analysis for p62 and PINK1 normalized to β-actin (related to Figure 2k) from MEFs and AEFs mock or MGT retroviral-transduced at D12. (h) Representative flow cytometry scatter plots of cardiac troponin (cTnT) in MEFs and AEFs mock or MGT retroviral-infected on D11, untreated or supplemented with rapamycin (10 nM) or urolithin A (UroA, 5 µM) (treatment from D4 to D11). Graphical data are mean ± s.d. of n = 10/43 (a-c) cells analyzed from two independent experiments, n=3 biological replicates from one representative experiment (e) or n = 4-5 biological replicates from two representative experiments (f, g). Each point on the plot (a-c, e-g) represents individual measurements. p values were calculated by Kruskal Wallis with false discovery rate correction and one-way ANOVA (a-c) and Tukey’s post-test for multiple comparisons (e, f, g). **p < 0.01; ****p < 0.0001 between the indicated groups. Graphs were created using GraphPad Prism (RRID:SCR_002798) and flow cytometry plots (h) with FlowJo (RRID:SCR_008520).

FIGURE S3. Metabolic modulation can bypass epigenetic and age-associated barriers to DCC. (a-d) Representative confocal microscopy images of α-MHC-GFP, mitochondria HSP-60 and DAPI in icMEFs with no Dox or Dox for 3 days (a). Analysis of TOM20 fluorescence corrected by cell area (b), form factor (shape, c) and branches per mitochondria (network connectivity, d) in icMEFs with no Dox or Dox (GFP-expressing cells) for 3 days. (e-f) Flow cytometry quantification of CellROX (ROS) (e) and mitochondrial mass by MitoTracker deep red staining f) depicted as the fold change of the relative median fluorescence intensity (MFI) in icMEFs with no Dox or Dox for 3 days and in the presence of Resveratrol (RVT, 20 nM). (g-m) Mass spectrometry-based histone acetylation and methylation proteomic bulk analysis in neonatal mouse cardiac myocytes (NMCM) and fibroblasts (NMCF), MEFs and AEFs MGT retroviral-infected at D11 in the presence of standard (S: glucose 25 mM, FBS 10%), supplementation with sodium acetate and α-KG (S*: sodium acetate 5 mM and α-KG 1,5 mM), low glucose (LG: glucose 1 mM, FBS 10%) or low lipids (LL: glucose 25 mM, FBS 1%) growth media formulation. PCA based on histone PTM data obtained from the z scores of the samples shown in the heatmap display of histone PTM levels (g). L/H (light/heavy) relative abundances ratios were obtained using a spike-in strategy (light channel: sample, heavy channel: spike-in standard), and were normalized over the average ratios across samples (h). Histone peptides were clustered based on Pearson’s correlation. The grey color indicates peptides that were not quantified. The panel on the right shows significant changes between NMCM and the indicated comparisons. Display of selected histone modified peptides showing the levels of mono-methylated form of H3 lysine 9 and mono-acetylated lysine 14 (i) and unmodified form of H3 lysine 4 (j) in MEFs samples, NMCM and NMCF and bi-acetylated form of H4 4–17 (k), mono-methylated form of H3 lysine 9 and mono-acetylated lysine 14 (l) and mono-methylated form of H3 lysine 9 (m) in AEFs samples, NMCM and NMCF, the bar indicates a p-value<0.05. Graphical data are mean ± s.d. of n = 9-10 cells analyzed (b-d) or n = 3-5 biological replicates (e-m) from two independent or one representative experiment. Each point on the plot (b-g) indicates individual measurements and means (i-m). p values were calculated by one-way ANOVA or Kruskal Wallis with False Discovery Rate correction by post-hoc Tukey’s test. *p < 0.05; **p < 0.01 with respect to No Dox or between the indicated groups. Data analysis and visualization were performed using Perseus (RRID:SCR_015753) and GraphPad Prism (RRID:SCR_002798). Scale bar, 20 µm in (a).

FIGURE S4. Metabolic modulation can bypass chromatin and aging-associated barriers to direct cardiac conversion. (a) Body weight (grams, g) of animals under control (CD), high fat (HFD) and low fat/high fructose (LFD) dietary regimens from week 0, during 10 weeks of diet intervention. (b) Serum glucose serum concentration (mg/dL, fold to control diet, CD) from CD, HFD and LFD animals at the end of the dietary regimen. (c-d) Densitometric quantification of immunoblotting analysis for total histone H3 (c) and H3 pan-acetylated (acetyl K9 + K14 + K18 + K23 + K27) (d) normalized to Ponceau S staining from whole cell extracts of CFs isolated from animals under control diet (CD), high fat (HFD) and low fat/high fructose (LFD) at the end of the dietary regimen. (e) qPCR analysis of the relative expression of the indicated genes in AEFs isolated from animals under control diet (CD), high fat (HFD) and low fat/high fructose (LFD) at the end of the dietary regimen and subjected to mock or MGT retroviral-transductions and analyzed on D11/12. (f) Confocal microscopy analysis of TMRE live cell imaging and quantification depicted as relative mean fluorescence intensity in AEFs isolated from animals under control diet (CD), high fat (HFD) and low fat/high fructose (LFD) at the end of the dietary regimen and subjected to mock or MGT retroviral-transductions and analyzed at D11/12. Graphical data are mean ± s.e.m of n = 7-9 mice (a, b), n = 4-6 biological replicates (c, d, f) from two independent experiments or n = 3 biological replicates from one representative experiment (e). Each point on the plot indicates individual measurements (b, c, d, f). p values were calculated by one-way ANOVA with False Discovery Rate followed by Tukey’s post-test for multiple comparisons. *p < 0.05; **p < 0.01; ***p < 0.001 with respect to HFD (a) or the indicated groups. Graphs were created using GraphPad Prism (RRID:SCR_002798).

FIGURE S5. Uncropped membranes relative to the immunoblotting for PGC-1α, p62, PINK1 and β-actin displayed in Figure S3F and Figure 3K and total Histone H3, H3 pan-acetylated (acetyl K9 + K14 + K18 + K23 + K27) or Ponceau S staining displayed in Figure 5d.

**Table S1.** Mass spectrometry-based histone acetylation and methylation proteomic bulk analysis in mock and MGT-transduced MEFs and AEFs at D11, and HL-1 mouse cardiomyocyte cell line. Light/Heavy (L/H) ratios of % relative abundance (%RA) value for the indicated samples.

**Table S2.** LC-MS-based untargeted metabolomics of MEFs mock- (control) or MGT-transduced at D11, and the HL-1 mouse cardiomyocyte cell line. List of 99 validated metabolic species identified and relative abundance.

**Table S3.** LC-MS-based untargeted metabolomics of MEFs mock- (control) or MGT-transduced at D11, and the HL-1 mouse cardiomyocyte cell line. Differential analysis of cellular metabolites was associated with top 18 significantly altered metabolic pathways, in terms of -log(p value) by FDR.

| **Pathway** | **p-Value** | **Metabolites** | **FDR** | **-Log(p)** | **Impact** |
| --- | --- | --- | --- | --- | --- |
| Aminoacyl-tRNA biosynthesis | 1.8613e-9 | 14/48 | 1.5635e-7 | 8.7302 | 0.0 |
| Alanine, aspartate and glutamate metabolism | 9.4806e-5 | 7/28 | 0.0039819 | 4.0232 | 0.621 |
| Arginine biosynthesis | 1.6352e-4 | 5/14 | 0.0045785 | 3.7864 | 0.19289 |
| Arginine and proline metabolism | 7.2478e-4 | 7/38 | 0.012954 | 3.1398 | 0.37647 |
| Glutathione metabolism | 7.7104e-4 | 6/28 | 0.012954 | 3.1129 | 0.06811 |
| Phenylalanine metabolism | 0.0010706 | 4/12 | 0.014767 | 2.9704 | 0.59524 |
| Purine metabolism | 0.0012306 | 9/66 | 0.014767 | 2.9099 | 0.26679 |
| Nicotinate and nicotinamide metabolism | 0.0026847 | 4/15 | 0.028189 | 2.5711 | 0.56711 |
| Valine, leucine and isoleucine biosynthesis | 0.0033639 | 3/8 | 0.031396 | 2.4732 | 0.0 |
| Pyrimidine metabolism | 0.0046484 | 6/39 | 0.039047 | 2.3327 | 0.15316 |
| beta-Alanine metabolism | 0.0097417 | 4/21 | 0.068525 | 2.0114 | 0.05597 |
| Phenylalanine, tyrosine and tryptophan biosynthesis | 0.0097892 | 2/4 | 0.068525 | 2.0093 | 1.0 |
| Glycerophospholipid metabolism | 0.015089 | 5/36 | 0.097499 | 1.8213 | 0.10756 |
| Nitrogen metabolism | 0.023177 | 2/6 | 0.12979 | 1.6349 | 0.0 |
| D-Glutamine and D-glutamate metabolism | 0.023177 | 2/6 | 0.12979 | 1.6349 | 0.5 |
| Histidine Metabolism | 0.026466 | 3/16 | 0.13895 | 1.5773 | 0.22131 |
| Taurine and hypotaurine metabolism | 0.040987 | 2/8 | 0.20252 | 1.3874 | 0.71428 |
| TCA cycle | 0.047852 | 3/20 | 0.22331 | 1.3201 | 0.12311 |

**Table S4.** Mass spectrometry-based histone acetylation and methylation proteomic bulk analysis in neonatal mouse cardiac myocytes (NMCM) and fibroblasts (NMCF), MEFs and AEFs MGT retroviral-infected at D11 in the presence of standard (S: glucose 25 mM, FBS 10%), supplementation with sodium acetate and α-KG (S*: sodium acetate 5 mM and α-KG 1,5 mM), low glucose (LG: glucose 1 mM, FBS 10%) or low lipids (LL: glucose 25 mM, FBS 1%) growth media formulation. Light/Heavy (L/H) ratios of % relative abundance (%RA) value for the indicated samples.

**Table S5** – List of primers used for measurements of gene expression levels by RT- qPCR.

| Name | Sequence |
| --- | --- |
| m*18*S-F | 5’- AGGATGTGAAGGATGGGAAGT-3’ |
| m*18S-*R | 5’- CAGGTCCTCACGCAGCTTG-3’ |
| m*Mef2c*-F | 5’- GAGCCGGACAAACTCAGACA-3’ |
| m*Mef2c*-R | 5’- TCAAAGCTGGGAGGTGGAAC-3’ |
| m*Tnnt2*-F | 5’- GAGCTACAGACTCTGATCGAGG-3’ |
| m*Tnnt2*-R | 5’- CCGCTCATTGCGAATACGC-3’ |
| m*Col3a1*-F | 5’- CTGTAACATGGAAACTGGGGAAA-3’ |
| m*Col3a1*-R | 5’- CCATAGCTGAACTGAAAACCACC-3’ |
| m*Acta1-*F | 5’- GGATTCTGGCGATGGTGTAAC-3’ |
| m*Acta1-*R | 5’-GACCAGCTAGATCCAGGCG-3’ |
| m*Myl7-*F | 5’-TCGGGAGGGTAAGTGTTCC-3’ |
| m*Myl7-*R | 5’-GTCCGTCCCATTGAGCTTCTC-3’ |
| m*Tbx5-*F | 5’-CTTCTATCGCTCGGGCTACC-3’ |
| m*Tbx5-*R | 5’-GCTATAGGAGGGCATGCTGG-3’ |
| m*Col1a1*-F | 5’-GCTCCTCTTAGGGGCCACT-3’ |
| m*Col1a1*-R | 5’-CCACGTCTCACCATTGGGG-3’ |
| m*Eln*-F | 5’-TGTCCCACTGGGTTATCCCAT-3’ |
| m*Eln*-R | 5’-CAGCTACTCCATAGGGCAATTTC-3’ |
